# Kaposi’s sarcoma-associated herpesvirus forms and maintains R-loops at origins of lytic replication

**DOI:** 10.64898/2026.08.31.748444

**Authors:** Mariel Kleer, Julia Fox, Jennifer A. Corcoran

## Abstract

GC-rich sequences are abundant in human herpesviruses genomes. GC-rich regions can form three-stranded RNA:DNA hybrid structures called R-loops. Though these hybrid structures serve important biological roles at telomeres or during cellular DNA synthesis, unscheduled or prolonged R-loop formation causes DNA damage and genome instability. For this reason, several mechanisms exist to resolve R-loops including endoribonucleases RNaseH1 (constitutively expressed) and RNaseH2A (cell cycle-regulated) which degrade the RNA portion of the R-loop. The Kaposi’s sarcoma-associated herpesvirus (KSHV) origins of lytic replication (OriLyts) contain multiple *cis*-acting elements that are required for viral DNA replication including the production of GC-rich and repetitive transcripts, *T1.4* (OriLyt-L) and *kaposin* (OriLyt-R). We previously showed that R-loops form at both OriLyts and that deleting *kaposin* repeats or decreasing their GC-rich content prevented R-loop formation at OriLyt-R, reduced genome amplification after primary infection and caused defects in latency establishment. To define the contribution that R-loops play in KSHV replication, we overexpressed RNaseH1, reasoning that excess RNaseH1 would resolve both OriLyt R-loops. However, RNaseH1 protein levels decreased following KSHV reactivation in both iSLK and BCBL-1 cell lines. Using co-transfection, we discovered that the KSHV viral replication and transcription activator protein, RTA, mediated RNaseH1 protein decreases in a E3 ligase domain-dependent manner without impacting levels of its cognate RNA transcript. We attempted to construct an RTA-resistant yet functional version of RNaseH1 by site-directed mutagenesis of lysine residues individually or in combination, yet these constructs remain susceptible to RTA-mediated protein decreases. An amino terminally tagged RNaseH1 displayed reduced susceptibility to RTA, suggesting that RTA may target the N-terminus of RNaseH1 for ubiquitination. However, overexpression of the cell-cycle regulated endonuclease, RNaseH2, exhibited RTA resistance, suggesting RNaseH2 may be a tool that will effectively resolve R-loops during KSHV infection. KSHV is not the only herpesvirus to encode a protein that reduces RNaseH1 levels, as co-expression of RTA homologs from the related γ-herpesviruses EBV and MHV-68 likewise decreased steady-state levels of RNaseH1 protein. We propose that RTA-mediated RNaseH1 degradation is conserved strategy to ensure R-loop persistence during γ-herpesvirus infection, underscoring the importance of these structures.

## INTRODUCTION

Kaposi’s sarcoma-associated herpesvirus (KSHV) is the causative agent of the endothelial neoplasm, Kaposi’s sarcoma (KS), and the lymphoproliferative disorders, primary effusion lymphoma (PEL) and multicentric Castleman’s disease (MCD)^1,2^. The prevalence of KSHV varies geographically; seropositivity is highest in Sub-Saharan Africa (>40-90%), between 20-30% in the Mediterranean, and <10% in Europe, Asia, and North American^3^. The KSHV dsDNA genome is ∼170 kbp in size with a 140 kbp unique coding region that is flanked by a 30 kbp terminal repeat (TR) region. The TR region is composed of 801 bp repeats that are highly GC-rich and contains the latent origin of replication (OriP). The KSHV genome also contains two origins of lytic replication (OriLyts) termed OriLyt-L(left) and OriLyt-R(right) which likewise contain GC-rich and repetitive elements. The unique coding region encodes approximately 80 open reading frames (ORFs), 25 mature miRNAs and as at least five long non-coding RNAs (lncRNAs)^4^. Like other herpesviruses, KSHV adopts a biphasic replication cycle comprised of lytic reactivation, and a dormant phase called viral latency.

Following the infection of most cell types, KSHV establishes latency within 48-72 hours^5,6^. KSHV latency has been studied extensively; however, the events which precede its establishment represent a unique and poorly understood phase of viral replication known as primary infection. This period represents a critical phase of KSHV infection as a failure to properly establish latency translates to a failure to successfully reactivate, leading to a dead-end for viral replication. The goals of primary infection are to i) evade the host immune response ii) increase viral genome copy number and iii) to establish proper chromatinization, tethering, and organization of the viral genome to facilitate future reactivation. Following entry, the KSHV genome undergoes a bi-phasic euchromatin to heterochromatin transition where initial activating histone marks such as H3K4me3 and H3K27ac are first deposited on viral DNA. These marks correlate with a burst of viral gene transcription that peaks at two to eight hours post-infection^5–7^. This transient phase of gene expression includes viral genes required for boosting subsequent viral transcription such as replication and transcription activator (RTA), those involved in modulating immune responses and those involved in DNA replication such as the viral processivity factor (ORF59) and the viral DNA polymerase (ORF9). The repertoire of proteins expressed immediately post primary infection led to the hypothesis that this brief burst of gene expression serves to combat the host immune response and amplify the incoming viral genome to increase genome copy number^7^. In support of this, treatment with the viral DNA polymerase inhibitor, phosphonoacetic acid (PAA), results in a decreased latent viral genome copy number. Likewise, primary infection with a recombinant KSHV virus deficient in ORF59 expression, which is required for active DNA replication, also results in a failure to increase viral genome copy number following primary infection^7^. These observations suggest that genome amplification after primary infection proceeds in a similar fashion to replication during lytic reactivation but is halted prior to virion production culminating instead in latency establishment.

After reactivation, the requirements for genome amplification have been well described. KSHV lytic DNA replication is initiated at one of two possible origins, origin of lytic replication L (OriLyt-L) or origin of lytic replication R (OriLyt-R). These duplicate KSHV OriLyts contain similar *cis*-acting sequences in reverse orientation from one another (Figure 1). Yan Yuan’s lab demonstrated four *cis* sequences are absolutely required for genome replication (reviewed in^8^). These include i) an 18 bp AT-rich palindromic sequence, believed to be required for DNA unwinding, ii) eight C/EBP binding motifs within a 240 bp region which enable host C/EBP binding, iii) an RRE, which is required both for the transcription of a GC-rich RNA (*T1.4* for OriLyt-L and *kaposin* for OriLytR) and the recruitment of viral replication machinery, and iv) the GC-rich repeat sequences, the function of which is unclear^9^. Initial work characterizing the KSHV OriLyts revealed that, in Vero cells, OriLyt-L is required for viral genome replication whereas OriLyt-R is not^10^. This study led the field to largely disregard the role OriLyt-R in favour of OriLyt-L. Yet work performed using the related herpesvirus, murine gammaherpesvirus 68 (MHV-68), showed the OriLyts can be utilized in a cell type specific manner as each is bound by a distinct subset of cellular proteins^11^. This suggests that while OriLyt-L may be required for KSHV DNA replication in Vero cells, that OriLyt-R may be required for DNA replication in other cell types or during other replication phases. Lytic KSHV DNA replication also requires a total of eight *trans*-acting viral proteins. The first proteins to be recruited to the OriLyts are RTA and K8α. RTA is recruited by binding of the RRE contained within the T1.4 (OriLyt-L) or kaposin (OriLyt-R) promoters, whereas K8α is thought to be recruited either by engaging with C/EBP or by binding the OriLyt-associated RNAs, *T1.4* or *kaposin*^9,12,13^. RTA and K8α fulfill the roles of viral origin binding proteins (OBPs) and associate with the six core herpesvirus replication proteins which include the single stranded DNA binding protein (ssDBP, [ORF6]), viral DNA polymerase

**Figure 1.**
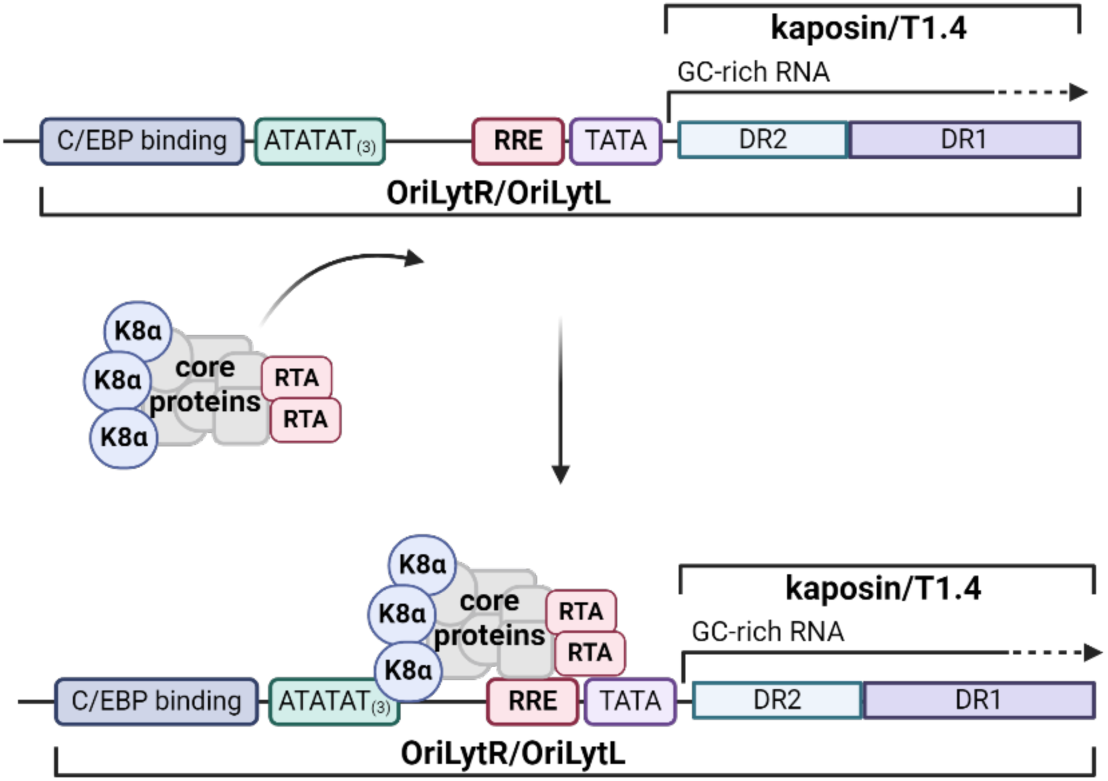
Lytic DNA replication from the KSHV OriLyts. The major *cis* features of the KSHV OriLyts are shown including the C/EBP binding site, the AT-rich region, the RTA-responsive element, and the TATA box and GC-rich associated transcripts T1.4 and kaposin for OriLyt-L and OriLyt-R respectively. The *trans* acting factors RTA, K8α, and the six core herpesvirus replication proteins ORF6, ORF9, ORF40/41, ORF44, ORF56 and ORF59 are shown being recruited to the OriLyt, following which replication can be initiated. Adapted from^8^.

(POL, [ORF9]), primase-associated factor (PAF, [ORF40/41]), helicase (HEL, [ORF44]), primase (PRI, [ORF56]) and polymerase processivity factor (PPF, [ORF59])^8^. Together, these eight viral proteins form the viral replication initiation complex and the loss of any these proteins abolishes DNA replication from an OriLyt containing plasmid^14^. Many cellular proteins are also recruited to the viral replication initiation complex and some, such as topoisomerase 1 and 2, are required for viral DNA replication^15^.

Protein degradation is an important mechanism that allows cells to fine-tune protein concentrations and eliminate those proteins which have been improperly folded. Proteins can be degraded through autophagy but are more commonly degraded through the ubiquitin-proteasome system (reviewed in^16,17^). Ubiquitin is a small 76-amino acid modifier that can be reversibly added to proteins as a post-translation modification (PTM). Ubiquitin is primarily added to internal lysine residues, although it can also be added to cysteine, serine, and threonine residues, as well as the N-termini of proteins^18–20^. Ubiquitin is added via a series of sequential reactions by activating (E1), conjugating (E2), and ligating (E3) enzymes (reviewed in^21^). The specificity of the ubiquitin system is determined by the interaction of E3 enzymes, of which there are over 600^22^, with their target proteins. E3 ligases can either directly (Homologous to E6AP C-terminus (HECT) and RING-in-between-RING (RBR) E3 ligases) or indirectly (Really Interesting New Gene (RING) and U-box E3 ligases) transfer ubiquitin to the target protein. An extraordinary diversity of ubiquitin linkages can be produced to signal a wide variety of outcomes. The most well characterized ubiquitin linkage is Lys48 (homogenous polyubiquitination), which serves to target proteins to the proteasome where they are recognized, unfolded, and degraded into smaller peptides.

Viruses can encode their own E3 ligases to target cellular or viral proteins for degradation. These molecules often target proteins that are antiviral in nature and when eliminated lead to increased viral replication. For example, HSV-1 ICP0 and rotavirus NSP1 are both viral RING or RING-like E3 ligases that promote the degradation of numerous cellular targets including classical antiviral molecules, such as MyD88 and IRF3, which would otherwise elicit immune detection^23,24^. KSHV encodes multiple E3 ligases including K3, K5, and the viral lytic switch protein, Replication and Transcription Activator (RTA)^25,26^. RTA is classified as a RING-like E3 ligase based on the cysteine and histidine rich region contained within its N-terminus (Figure 2) and its ability to undergo autoubiquitination^27^. RTA also contains multiple small ubiquitin-like modifier (SUMO)-interacting motifs (SIMs) that enable it to specifically interact with and target SUMO-modified proteins. For this reason, RTA is classified as a viral SUMO-targeting ubiquitin ligase (STUbL)^28^. Numerous RTA targets have been identified including antiviral molecules such as myeloid differentiation primary response 88 (MyD88), interferon response factor 7 (IRF7), and signal transducer and activator of transcription 6 (STAT6)^29–33^. More recent work has attempted to define the full repertoire of RTA targets during KSHV reactivation^34^. Identifying cellular proteins targeted by viral E3 ligases can help pinpoint pathways that are critical to viral success and provides an opportunity to identify novel antiviral molecules.

**Figure 2.**
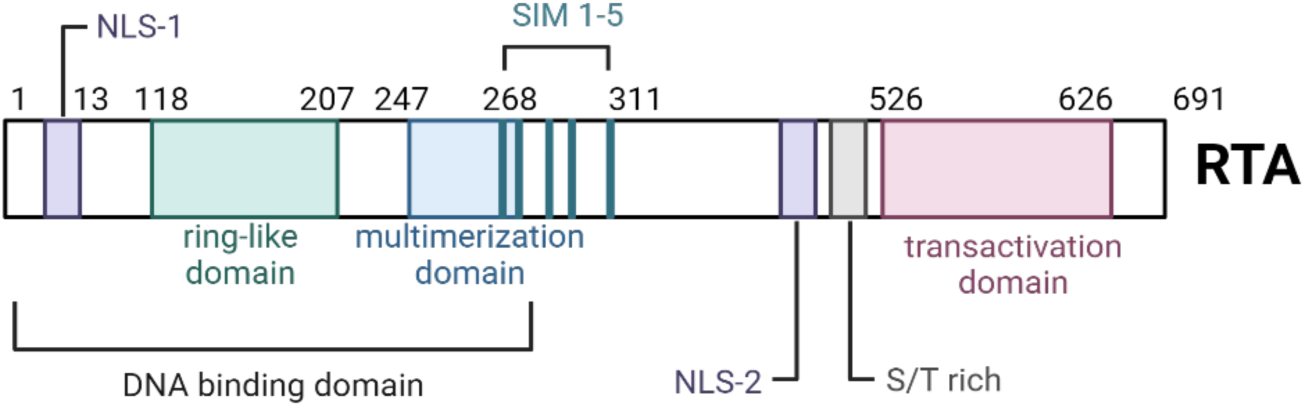
Functional domains of the replication and transcription transactivator RTA. A linear depiction of the position of RTA domains organized from left (amino acid 1) to right (amino acid 691). NLS; nuclear localization signal, SIM; sumo interaction motif, S/T-rich; serine/threonine rich.

R-loops are three-stranded nucleotide structures that most often form during transcription when the nascent transcript remains annealed to its template DNA strand, displacing the non-template DNA strand into a loop. R-loops were originally discovered *in vitro* in 1976, but were not confirmed to form in mammalian cells until 2003 when they were found be required for class switch recombination in B lymphocytes^35–37^. R-loops are thermodynamically stable and preferentially form in G-rich regions of DNA, particularly those with a strong GC-skew between the template and non-template strands^38,39^. R-loop formation aids many biological functions including transcription termination, telomere maintenance, promoter accessibility, bacteriophage and mitochondrial DNA replication, DNA repair, and genomic looping^40–49^. However, unscheduled or prolonged R-loop formation can lead to genome instability and so R-loop formation and resolution is tightly regulated by several factors^50–52^. These include proteins such as Topoisomerase I (Top1), which supresses R-loop formation by alleviating negative supercoiling^53^, RNA:DNA specific helicases such as senataxin (SETX) or Aquarius (AQR), which resolve R-loops by unwinding the and RNA:DNA specific endonucleases such as RNaseH1 or RNaseH2, which resolve R-loops by degrading the RNA portion of the hybrid^40,52–56^. Although both RNaseH1 and RNaseH2 function in R-loop removal, the two enzymes exhibit differential regulation. RNaseH1 is constitutively expressed at low levels independent of cell cycle and is a monomer, whereas RNaseH2 functions in a heterotrimeric protein complex and is expressed and chromatin-associated in a cell-cycle dependent manner^53^. Moreover, RNaseH2 is thought to account for the majority of cellular RNaseH activity, whereas RNaseH1 may bind chromatin but only appears to be activated in response to stress-induced high R-loop loads^53,57^. The decreased expression of any of these R-loop resolvers disrupts R-loop homeostasis and leads to R-loop accumulation^58–60^.

Viral genomes form a variety of regulatory structures including stem loops, internal ribosomal entry sites, hairpins, and G-quadraplexes, but until recently little was known about the prevalence of R-loops in viral genomes^61,62^. In 2011, Rennekamp et al. found that transcription of a GC-rich transcript at the two EBV viral origins of lytic replication (OriLyts) led to the formation of R-loops which are required for effective viral DNA replication^63^. In 2020, Tai-Schmiedel et al. found that HCMV also uses a GC-rich transcript to form an R-loop at its lone OriLyt and that, again, the formation of this structure is required for viral DNA replication^64^. In both cases, the authors showed that R-loop formation was required to express (HCMV) the viral ssDNA binding protein or recruit it to the OriLyt(s) (EBV), events that must precede viral genome replication. More recently, extensive computational analysis by Wongsurawat et al. revealed that R-loops are predicted to form in thousands of viral genomes and that herpesvirus genomes are particularly enriched for these structures, likely due to their high GC-rich genomic content^65^. The authors of this study validated their *in-silico* predictions by experimentally confirming R-loop formation in the terminal repeat (TR) and viral ORF16 promoter regions of the KSHV genome^65^. These data suggest that R-loop formation is a conserved feature of herpesvirus genomes.

*Kaposin*, the GC-rich transcript from OriLyt-R, is the most abundantly expressed viral RNA in KS tumors and one of the most abundantly expressed viral RNAs during every stage of KSHV infection^4,7,66–69^. *Kaposin* is an unusual RNA comprised largely of two sets of GC-rich direct repeats (DR1 and DR2) and is translated to produce at least three protein products: KapA, KapB, and KapC. The function of these products during KSHV replication has been of great interest but the GC-rich, repetitive, and polycistronic nature of *kaposin* has made it difficult to study. To address this, our lab generated a panel of kaposin recombinant KSHV viruses (ΔKapABC, ΔKapBC, ΔKapB, ΔKapC) based on the BAC16 system for the analysis of *kaposin* function during KSHV infection^70,71^. We intended to use this panel to analyze the role of the KapA, KapB, and KapC during KSHV infection; however, to our surprise, we found that all kaposin recombinant viruses exhibited decreased genome copy number relative to the wildtype (WT) BAC16 virus during both latency and following primary infection, regardless of which protein was deleted. We attempted to complement the defects we observed during BAC16ΔKapB virus infection by supplying KapB protein *in trans*, however we were unable to do so successfully. Revisiting our mutagenesis design, we realized that in every case where we had deleted a Kap protein, we had also drastically altered the *kaposin* RNA sequence^70^. These data led us to hypothesize a role for the *kaposin* transcript *in cis*. When expressed *in cis* from its genomic location, *kaposin* forms part of OriLyt-R (Figure 1). In each of our recombinant viruses, the *kaposin* transcript is still located within the OriLyt-R; however, the GC-rich region is either removed, making the transcript much shorter, or recoded, resulting in a transcript that is no longer repetitive or GC-rich. The other OriLyt-associated transcript, *T1.4*, likewise contains a GC-rich region that is required for viral DNA replication, and *T1.4* transcription is required for genome amplification during *de novo* KSHV infection^9,13^. We previously showed that both GC-rich transcripts *kaposin* and *T1.4 in cis* formed R-loops at the KSHV OriLyts; however, kaposin recombinant viruses lacking the GC-rich region in *kaposin* no longer form R-loops at OriLyt-R^72^. If R-loop formation is required for KSHV DNA replication, particularly the burst of amplification after primary infection, this could explain why kaposin recombinant viruses, which lack a GC-rich *kaposin* transcript and do not form an R-loop at OriLyt-R display a lower genome copy number relative to WT following primary infection. This hypothesis is supported by data from related herpesviruses, HCMV and EBV, which have been shown to use an RNA transcript in cis to form an R-loop at their respective OriLyts, the formation of which is required for successful viral DNA replication^63,64^.

## RESULTS

### Altered recruitment of viral proteins to OriLyts in the absence of R-loop formation

There are eight viral proteins recruited to the KSHV OriLyts during active viral DNA replication: RTA, K8α, and six core herpesvirus replication proteins which include the single stranded DNA binding protein (ssDBP [ORF6]), viral DNA polymerase (ORF9), primase-associated factor (ORF40/41), helicase (ORF44), primase (ORF56) and polymerase processivity factor (ORF59) (Figure 1)^8,73^. Having shown that R-loop formation occurs at both OriLyts, we asked whether the formation of this structure is required for the recruitment of the viral DNA replication machinery and subsequent viral genome copying^63,64^. Given the decreased intracellular genome copy number of kaposin recombinant viruses following primary infection, we predict R-loop mediated recruitment is particularly important at this stage of the viral lifecycle. However, the low efficiency of primary infection results in significantly less viral genomic material than latent or reactivated iSLK cell lines, makes it too difficult to perform large-scale assays such as Chromatin Immunoprecipitation qPCR (ChIP-qPCR) at this replication phase. For that reason, we chose to analyze the lytic reactivation phase of WT, ΔKapABC, and ΔKapB iSLK infection, reported to require the same viral proteins. During WT lytic reactivation, R-loops form at both OriLyt-L and OriLyt-R; however, during ΔKapABC and ΔKapB virus reactivation, R-loop formation at OriLyt-R is abolished^70^. If R-loop formation is required for viral replication protein recruitment, this means that recruitment should occur at both OriLyts during WT virus reactivation, but only at OriLyt-L during ΔKapABC and ΔKapB virus reactivation. To test this, we used ChIP-qPCR to assess RTA, K8α, and ORF6 recruitment to OriLyts during latency (negative control) and at 48 hours post-reactivation. To distinguish between regions of the OriLyts, we designed primers sets that were specific for either the gene body or the promoter sections of both T1.4 and kaposin. The RTA-responsive element (RRE) that RTA binds to directly is contained within the promoter region; therefore, we predicted we would observe RTA binding in the promoter and not in the gene body. In contrast, the precise binding sites for K8α and ORF6 are not known. As expected, we found there was no significant enrichment of RTA or K8α binding at either OriLyt during WT, ΔKapABC, and ΔKapB iSLK latency (Figure 3). ORF6 binding during latency was not tested due to a limited quantity of antibody. At 48 hours post WT KSHV reactivation, RTA was recruited to the promoter, but not the gene body, of both T1.4 and kaposin, with a slight, but not significant, enrichment at the T1.4 promoter region (Figure 3A). During WT KSHV reactivation we found that K8α was recruited to both the gene body and promoter of both OriLyts with similar efficiencies (Figure 3B). During ΔKapABC and ΔKapB reactivation, we found that K8α was likewise recruited to both promoters and gene bodies but exhibited a non-statistically significant increase in K8α binding at the T1.4 gene body relative to kaposin. Finally, we found that ORF6 bound with similar efficiency to both the gene and promoter regions of kaposin and T1.4 during WT KSHV reactivation whereas during ΔKapB reactivation there was an increase in ORF6 binding to the T1.4 promoter versus the kaposin promoter (Figure 3C). Likewise, during ΔKapABC and ΔKapB reactivation, RTA also bound to the promoter regions, with a slight increase in enrichment at the OriLyt-L promoter (Figure 3A). During WT KSHV reactivation we found that K8α was recruited to both the gene body and promoter of both OriLyts with similar efficiencies (Figure 3B).

**Figure 3.**
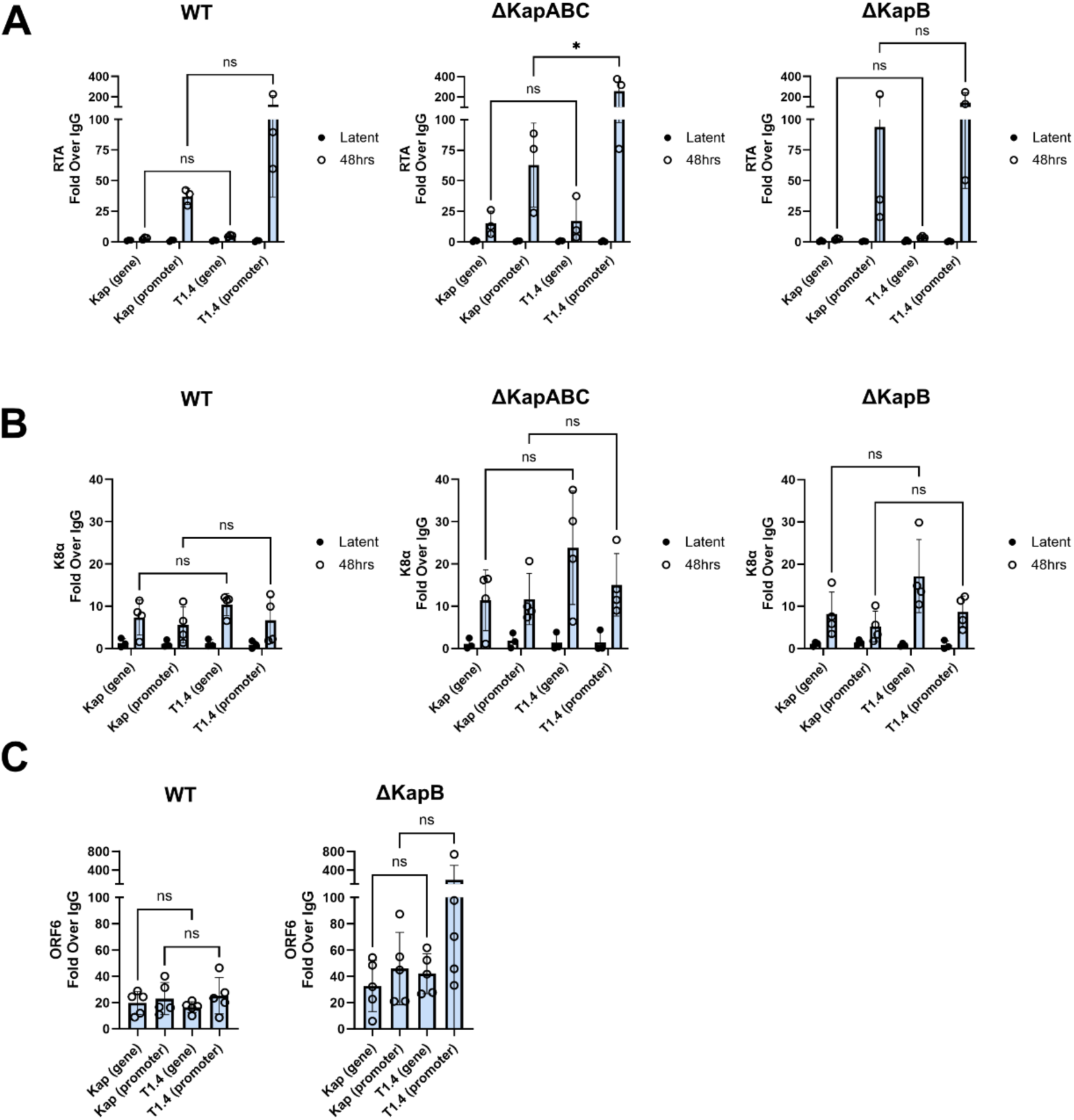
Altered recruitment of viral proteins to OriLytR/kaposin in the absence of R-loop formation. (A-B) WT, ΔKapABC and ΔKapB iSLK cells were seeded and either untreated (latency) or treated with Dox for 48 hrs to induce lytic reactivation. Cells were then crosslinked and ChIP-qPCR was performed using either an RTA (A) or K8α (B) specific antibody and primers specific for either the promoter or gene body regions of T1.4 and kaposin. Data is presented as fold change relative to the IgG control. Statistics were performed using a mixed-effects analysis with a Tukey’s post-hoc test. N≥3; mean ± SD. (C) WT and ΔKapB iSLK cells were seeded and treated with Dox for 48 hrs to induce lytic reactivation. Cells were then crosslinked and ChIP-qPCR was performed using an ORF6 specific antibody and primers specific for either the promoter or gene body region of T1.4 and kaposin. Data is presented as fold change relative to the IgG control. Statistics were performed using a repeated measures one-way ANOVA with a Tukey’s post host test. N=5; mean ± SD.

During ΔKapABC and ΔKapB reactivation, we found that K8α was likewise recruited to both promoters and gene bodies but exhibited a non-statistically significant increase in K8α binding at the T1.4 gene body relative to kaposin. Finally, we found that ORF6 bound with similar efficiency to both the gene and promoter regions of kaposin and T1.4 during WT KSHV reactivation, whereas during ΔKapB reactivation there was an increase in ORF6 binding to the T1.4 promoter versus the kaposin promoter (Figure 3C); however, this difference was not statistically significant. Taken together, these data indicate: i) R-loop formation is not required for RTA recruitment, ii) R-loop formation may promote K8α binding to the gene body, in agreement with the previous published role of *T1.4*-mediated K8α recruitment to OriLyt-L^13^, and iii) R-loop formation may promote ORF6 binding, in agreement with our knowledge of the role of R-loops is related herpesviruses^63^. However, given the large variance between experiments, further work is required to confirm these hypotheses.

### RNaseH1 protein levels decrease during KSHV reactivation

The absence of R-loop formation at the EBV and HCMV OriLyts corresponds to decreased viral genome replication^63,64^. Based on our findings of decreased genome copy number during kaposin recombinant virus latency and primary infection, we hypothesized R-loop formation may likewise be required for KSHV viral DNA replication. We previously showed that kaposin recombinant viruses can still undergo genome copying during lytic reactivation; however, an easy explanation for this is that the remaining OriLyt-L R-loop compensates for the loss of R-loop formation at OriLyt-R, allowing viral DNA replication to still occur. This, coupled with our inconclusive observations about the role of R-loops in viral replication protein recruitment, led us to seek to resolve R-loops at both OriLyts to better ascertain their role in viral replication. To achieve this, we chose to overexpress the cellular R-loop resolver, the RNA:DNA specific endonuclease, RNaseH1. RNaseH1-expressing lentiviruses were used to transduce latent WT iSLK-BAC16 cells and RNaseH1 expression was confirmed by immunoblotting for its C-terminal tag, V5 (Figure 4A).

**Figure 4.**
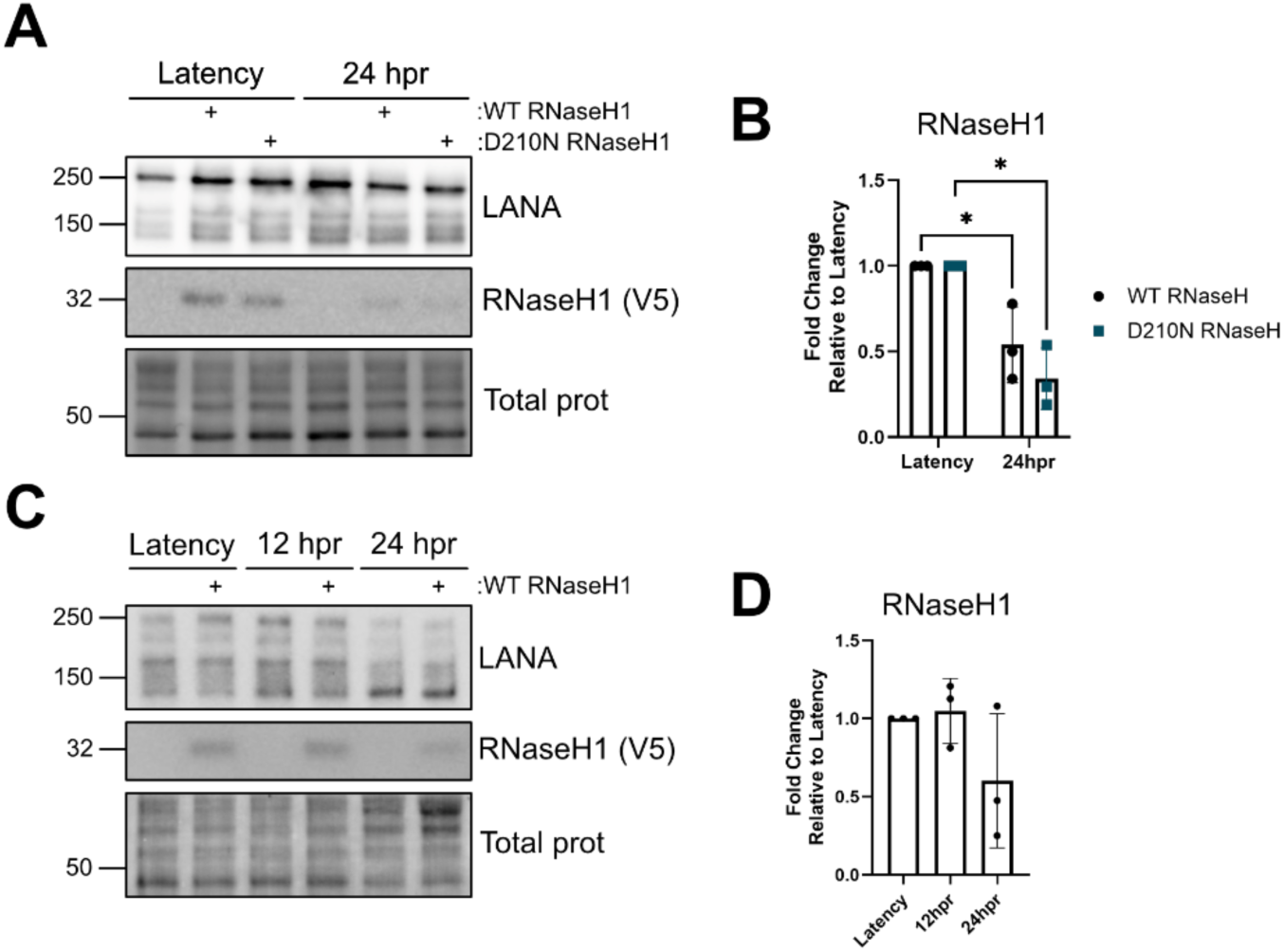
RNaseH1 protein expression level is decreased during KSHV reactivation. (A-D) iSLK (A-B) and BCBL (C-D) cells were transduced to express either WT RNaseH1, catalytically dead RNaseH1 (D210N RNaseH1), or an empty vector (EV) control. Cells were then either untreated (latency) or treated with Dox for 24 hrs (A-B) or 12 and 24 hrs (C-D) prior to lysis. Immunoblotting was performed using LANA and V5 (RNaseH1) specific antibodies. Samples were quantified by normalizing RNaseH1 protein levels to the total protein (total prot, explained in section 3.6.9) in each lane using ImageLab (BioRad) and then to the respective latency control. Statistics were performed using a two-way (B) or one-way (D) ANOVA with a Tukey’s post hoc test. N=3; mean ± SD. For panels A and C, one representative experiment of three is shown.

To test the effect of RNaseH1 overexpression on viral DNA replication, the transduced iSLKs were subsequently reactivated to enter the lytic lifecycle where active viral DNA replication from the OriLyts occurs. However, at 24 hours post-reactivation the steady-state level of RNaseH1 protein, as measured by immunoblotting, were noticeably decreased, rendering us unable to test our original hypothesis (Figure 4A,B). To test whether this effect was cell type specific, we repeated the same transduction experiment in a B-cell model of KSHV infection, BCBL-1s, and found that at 24 hours post-reactivation that RNaseH1 protein levels were again decreased relative to latency (Figure 4C,D). These data suggest that a viral gene product expressed early after lytic reactivation, but not latency, either decreases RNaseH1 expression or enhances the turnover of RNaseH1 mRNA or protein.

### RTA is sufficient for RNaseH protein decreases

The viral immediate early protein RTA is an essential viral transcription factor; however, RTA is also a sumo-targeting E3 ligase (STUbL) that target multiple host factors for degradation during infection^25,28–30^. For this reason, we tested whether RTA was sufficient to induce RNaseH1 levels to decrease during viral reactivation and determine whether this decrease occurred at the protein level, the RNA level, or both. Either WT RNaseH1, or a catalytically dead RNaseH1 (RNaseH1 D210N) were co-transfected with or without an RTA expression plasmid and RNaseH1 and RTA transcript levels measured 24 hours after transfection. Co-expression with RTA did not significantly alter RNaseH1 transcript levels (Figure 5A) but co-expression with both WT and catalytically inactive RNaseH1 led to an increase in RTA transcript levels, albeit non-statistically significant (Figure 5A). These data suggest that the expression of one construct does not significantly alter the steady-state RNA levels of the other. We then tested if RTA was sufficient to decrease RNaseH1 protein by co-transfection followed by immunoblotting. Relative to the EV control, co-expression of RTA lead to a significant decrease in WT RNaseH1 protein levels (Figure 5B,C), and co-expression of WT RNaseH1 led to a slight increase in RTA protein levels (Figure 5B,D). We reasoned that RNaseH1 protein turnover was enhanced by RTA, thus, we might be able to recover RNaseH1 protein levels by inhibiting cellular degradation pathways. In accordance with this, four hours prior to lysis, transfected cells were treated with either the proteasomal inhibitor (MG132), the autophagy inhibitor (chloroquine, CQ), or DMSO alone (control). Treatment with both MG132 and CQ resulted in a slight recovery of RNaseH1 protein levels (Figure 5B,C). However, we observed that both treatments induced a significant enhancement of RTA protein level (Figure 5B,D). Because RTA is autoubiquitinated, decreased RTA turnover when ubiquitin-mediated degradation is inhibited was not unexpected^25,31^. This means we are unlikely to achieve a complete restoration of RNaseH1 protein levels using degradation pathway inhibitors, as these inhibitors also block the turnover of RTA, leading to its increased abundance in the experiment. We hypothesized that the catalytic activity of RNaseH1 may promote RTA-mediated targeting of RNaseH1 protein. To test this, we repeated the same co-expression experiment using RNaseH1 D210N. We found that co-expression of RTA with RNaseH1 D210N led to a decrease RNaseH1 D210N protein level, but RNaseH1 D210N no longer led to an increase in RTA protein levels (Figure 5E-G). The mechanism by which WT, but not catalytically dead, RNaseH1 increases RTA protein levels is unknown. These data reveal that RTA, in the absence of any other viral protein, is sufficient to induce RNaseH1 protein decreases.

**Figure 5.**
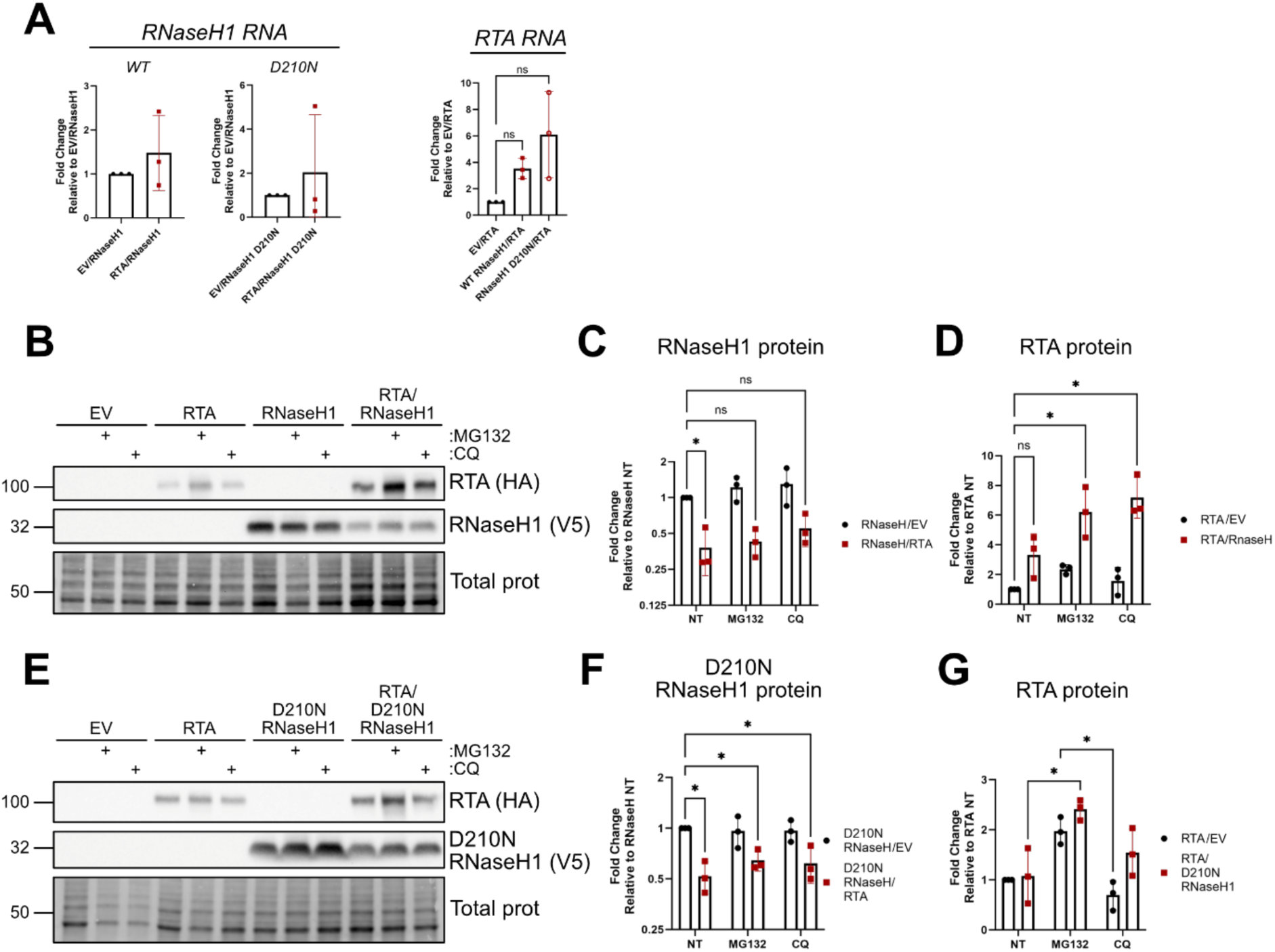
RTA is sufficient for RNaseH1 protein decrease. (A) HEK293T cells were transfected with an RNaseH1 (WT or D210N)-expressing vector in combination with an RTA-expressing vector or an empty vector (EV) control. 24 hrs post transfection RNA was extracted, and RT-qPCR was performed using *RNaseH1*, *RTA*, and *18S* (housekeeping) specific primers. Data is presented as fold change relative to the EV co-transfection control, aka RTA or RNaseH1 alone. Statistics were performed using a paired Student’s t-test (RNaseH1) or a one-way ANOVA with a Tukey’s post hoc test (RTA). N=3; mean ± SD. (B-G) HEK293T cells were transfected with an RNaseH1 (WT or D210N)-expressing vector or an EV control either alone or in combination with an RTA-expressing vector. 24 hrs post transfection cells were untreated (non-treated, NT) or treated with either MG132 (10 µM) or CQ (25 µM) for 4 hours prior to lysis. Immunoblotting was performed using RNaseH1 (V5) and RTA (HA) specific antibodies. Samples were quantified by normalizing RNaseH1 or RTA protein levels to the total protein (total prot) in each lane using ImageLab (BioRad) and then to the NT EV co-transfection control, aka RTA or RNaseH1 alone. Statistics were performed using a two-way ANOVA with a Dunnett’s post hoc test. N=3; mean ± SD. For both B and E, one representative experiment of three is shown.

### RTA E3 ligase activity and SIM domains are required for RNaseH1 protein decreases

RING-like E3 ligases coordinate two zinc ions that allow them to bind the E2 enzyme in a “cross-brace” position to facilitate the transfer of ubiquitin from the E2 to the target protein^27,74^. As an alternative strategy to test whether RTA targets RNaseH1 for degradation in a ubiquitin-specific manner, we used site-directed mutagenesis to create an E3 ligase-dead RTA construct by mutating a cysteine and a histidine residue within the RTA N-terminus as in^25,75^. The mutation of the cysteine and histidine residues within the E3 domain of RTA prevents the cross-bracing event that facilitates ubiquitin transfer; therefore, we will refer to this construct as E3 ligase dead RTA. If RTA-mediated ubiquitination of RNaseH1 is required to facilitate its degradation by the proteasome, co-expression of an E3 ligase-dead RTA will not be able catalyze the transfer of ubiquitin from the E2 enzyme to RNaseH1. If RNaseH1 is no longer ubiquitinated it should not be degraded and RNaseH1 levels during RTA co-expression should no longer be decreased. We co-transfected either WT or E3 ligase dead RTA with RNaseH1. The protein level of E3 ligase dead RTA was increased relative to WT RTA (Figure 6A), consistent with RTA auto-ubiquitination impairment when RTA E3 ligase activity was blocked. While co-expression of WT RTA decreased RNaseH1 levels, co-expression with E3 ligase dead RTA significantly restored RNaseH1 (Figure 6A,B). These data suggest that RTA E3 ligase activity is required for RNaseH1 turnover during co-expression. RTA contains several sumo interacting motif (SIM) domains that enable it to specifically target SUMO-modified proteins for degradation as a viral STUbL^28^. Although we could not find any information on the sumoylation status of RNaseH1 in the literature, we tested whether SUMO interaction is required for RTA-mediated RNaseH1 targeting. Using site-directed mutagenesis, we created a SIM-null RTA construct by mutating several amino acids within two of the five previously identified SIM domains, previously validated to destroy sumo-mediated RTA binding^28^. We co-transfected this construct, or WT RTA, with RNaseH1 in our co-transfection assay and measured RNaseH1 protein levels by immunoblotting. We found that co-expression with WT RTA led to a significant decrease in RNaseH1 protein levels, but co-expression with SIM-null RTA did not (Figure 6C,D). These data reveal that RTA requires SUMO-interacting motifs to decrease RNaseH1 protein levels.

**Figure 6.**
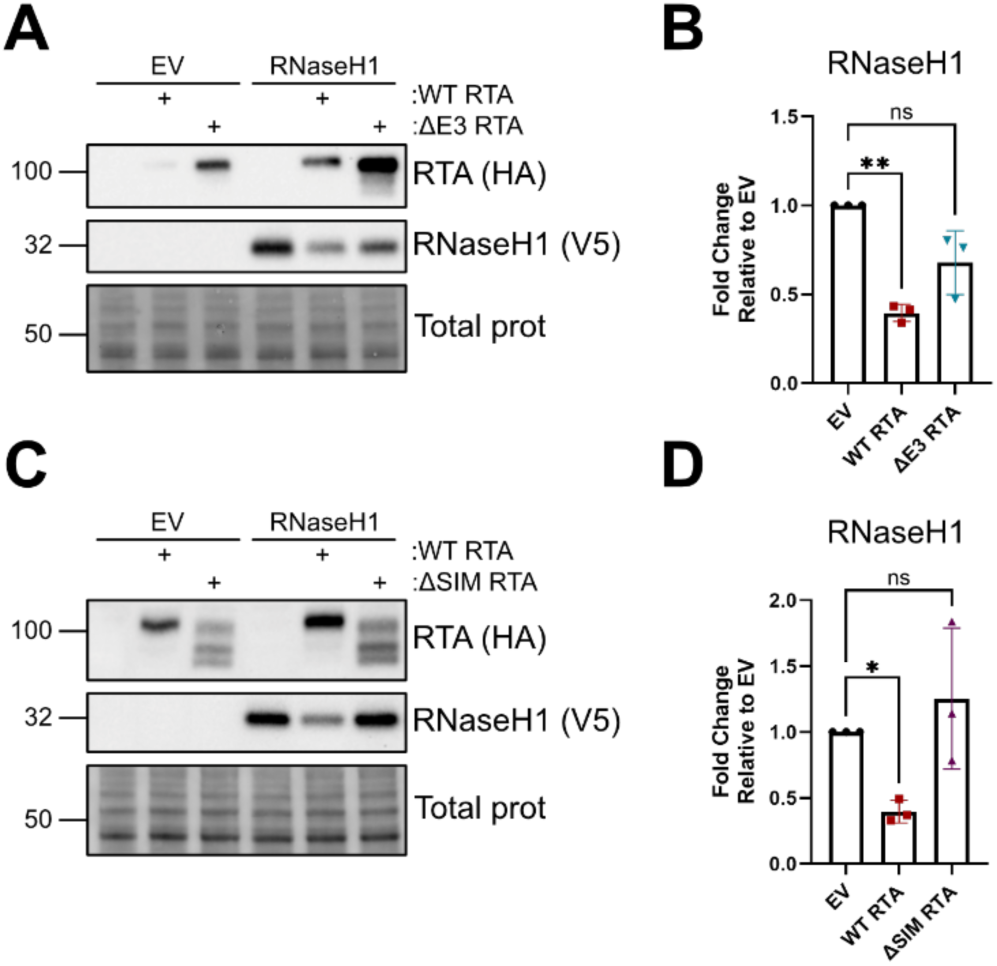
RTA STUbL activity is required to decrease RNaseH1 protein levels. (A-D) HEK293T cells were transfected with an RNaseH1-expressing vector in combination with an E3 ligase dead (A-B) or SIM-null (C-D) RTA-expressing vector or an empty vector (EV) control. 24 hrs post transfection cells were lysed and immunoblotting was performed using RNaseH1 (V5) and RTA (HA) specific antibodies. Samples were quantified by normalizing RNaseH1 protein levels to the total protein (total prot) in each lane using ImageLab (BioRad) and then to the respective EV control. Statistics were performed using a one-way ANOVA with a Dunnett’s post hoc test. N=3; mean ± SD. For panels A and C one representative experiment of three is shown.

RTA can either act as the E3 ligase that ubiquitinates a protein target directly or cooperate with cellular E3 ligases ITCH and UBE3C to enhance target protein ubiquitination^76,77^. To test if these proteins are required for RTA-induced RNaseH1 protein decreases, we designed shRNA sequences to silence ITCH and UBE3C transcripts and transduced cells with either ITCH- or UBE3C-targeting lentiviruses or a non-targeting (NT) control lentivirus. We confirmed the knockdown using RT-qPCR (Figure 7A,D) and then co-transfected RTA and RNaseH1 into NT control or ITCH- or UBE3C-silenced cells. By immunoblotting for RNaseH1 protein levels, we found that in the NT control cells RTA co-expression reliably corresponded to decreased RNaseH1 levels and that in both ITCH and UBE3C-silenced cells a similar decrease was observed (Figure 7B,C, E,F). From these data we conclude that ITCH and UBE3C are not required for RTA-mediated RNaseH1 turnover.

**Figure 7.**
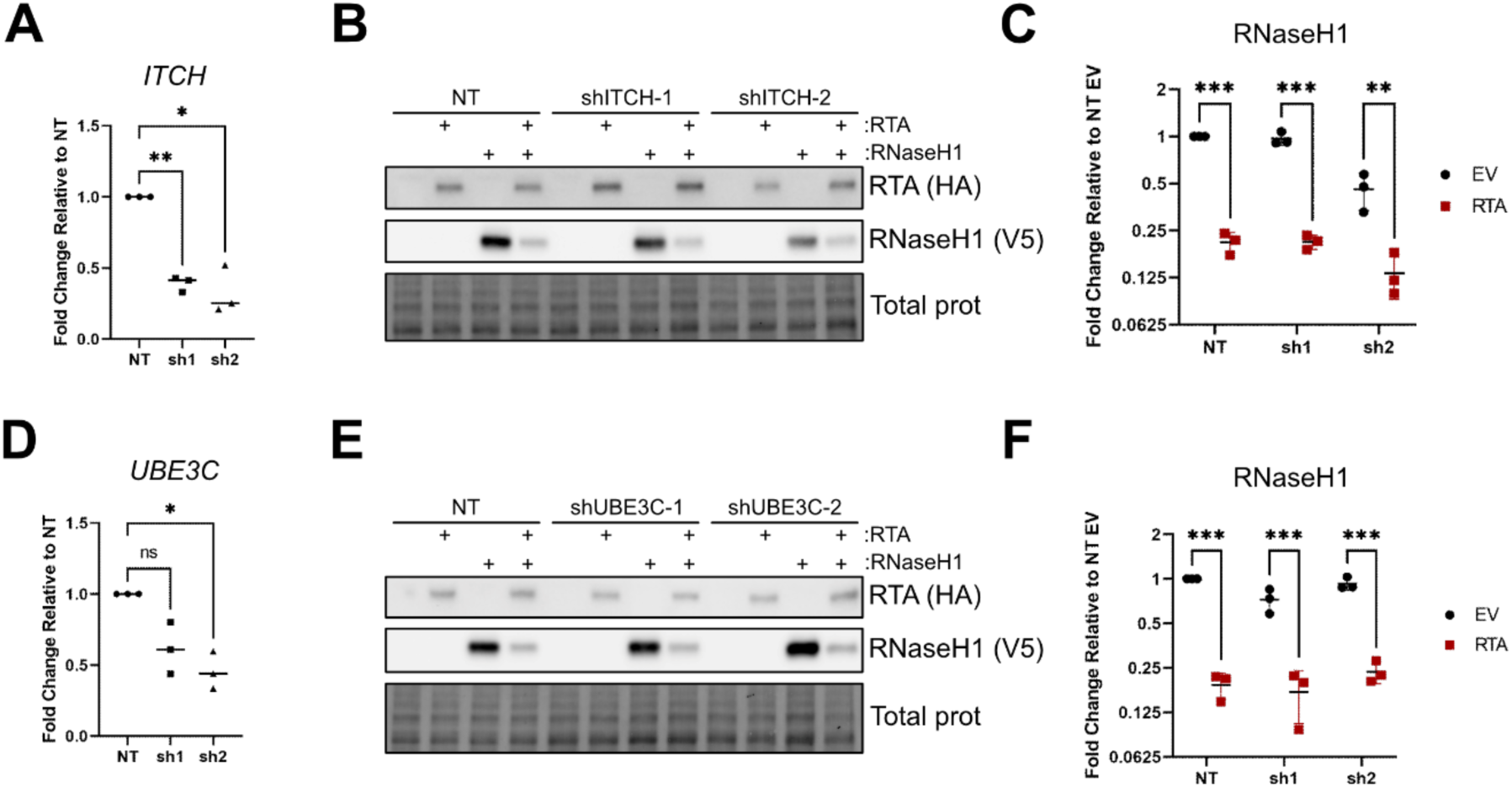
Host E3 ligases ITCH and UBE3C are not required for the RTA-mediated decrease in RNaseH1 protein. (A-F) HEK293T cells were transduced with lentiviruses expressing ITCH or UBE3C targeting shRNAs or a NT control. 24 hrs post transduction cells were either lysed to confirm knockdown (A, E) or transfected with an RNaseH1-expressing vector in combination with an RTA-expressing vector or an empty vector (EV) control (B-C, F-D). For lysed cells, RT-qPCR was performed using *ITCH*, *UBE3C*, or *18S* (housekeeping) specific primers. Data is presented as fold-change relative to NT. Statistics were performed using repeated-measure one-way ANOVA with a Dunnett’s post hoc test. N=3; mean ± SD. For transfected cells immunoblotting was performed using RNaseH1 (V5) and RTA (HA) specific antibodies. Samples were quantified by normalizing RNaseH1 protein levels to the total protein (total prot) in each lane using ImageLab (BioRad) and then to the respective EV control. Statistics were performed using a two-way ANOVA with a Tukey’s post hoc test. N=3; mean ± SD. For panels B and E one representative experiment of three is shown.

According to UniProt, RNaseH1 is predicted to be ubiquitinated at seven out of a total of 17 lysine residues: K60, K73, K119, K192, K198, K236, and K241 (Figure 8).

**Figure 8.**
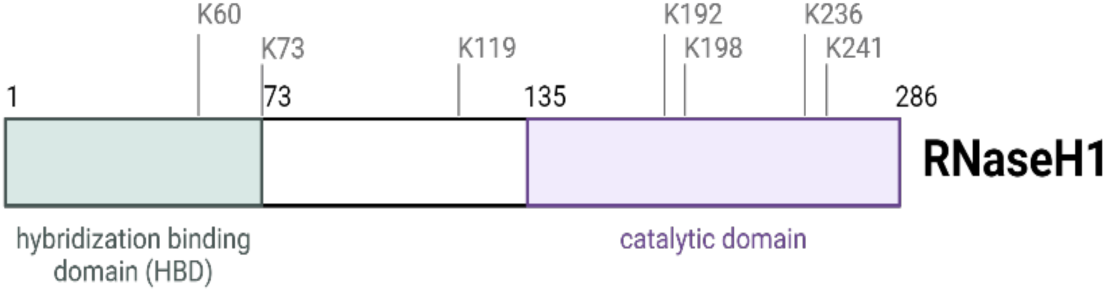
Functional domains of the RNA:DNA specific endonuclease RNaseH1. A linear depiction of the position of RNaseH1 domains organized from N-(left; amino acid 1) to C-terminus (right; amino acid 286). Lysine residues predicted to undergo ubiquitination are indicated.

Knowledge about the specific RNaseH1 lysine residue(s) targeted by RTA could allow us to create an RTA-resistant RNaseH1 construct with minimal effect on RNaseH1 function. To map which lysine residue is targeted by RTA we individually mutated these seven lysine residues to alanine. If RTA ubiquitinates a specific RNaseH1 lysine residue, then mutating this residue to alanine should render it resistant to this modification and block subsequent RNaseH1 degradation. We screened our set of K-mutant RNaseH1 constructs by co-transfection with or without RTA, performed immunoblotting, and quantified whether any singular K-to-A mutation was sufficient to prevent RTA-mediated RNaseH1 protein decreases. Some of these constructs expressed better than others and so in consideration of these expression differences, each mutant is graphed relative to its own EV co-expression control. We found that five out of seven RNaseH1 constructs (RNaseH1 K60A, K63A, K119A, K198A, and K236A) were significantly decreased during RTA co-expression (Figure 9A-D). Although two constructs (RNaseH1 K192A and K241A) no longer exhibited a statistically significant decrease in RNaseH1 levels, appears to be due to a single outlier experiment in both cases. To test whether any internal lysine residues could be used as ubiquitination targets, we created a lysine null (K-null) RNaseH1 construct by substituting all 17 lysine residues with alanine using site-directed mutagenesis. We co-transfected the RNaseH1 K-null construct with RTA or an EV control and performed immunoblotting to analyze its expression.

**Figure 9.**
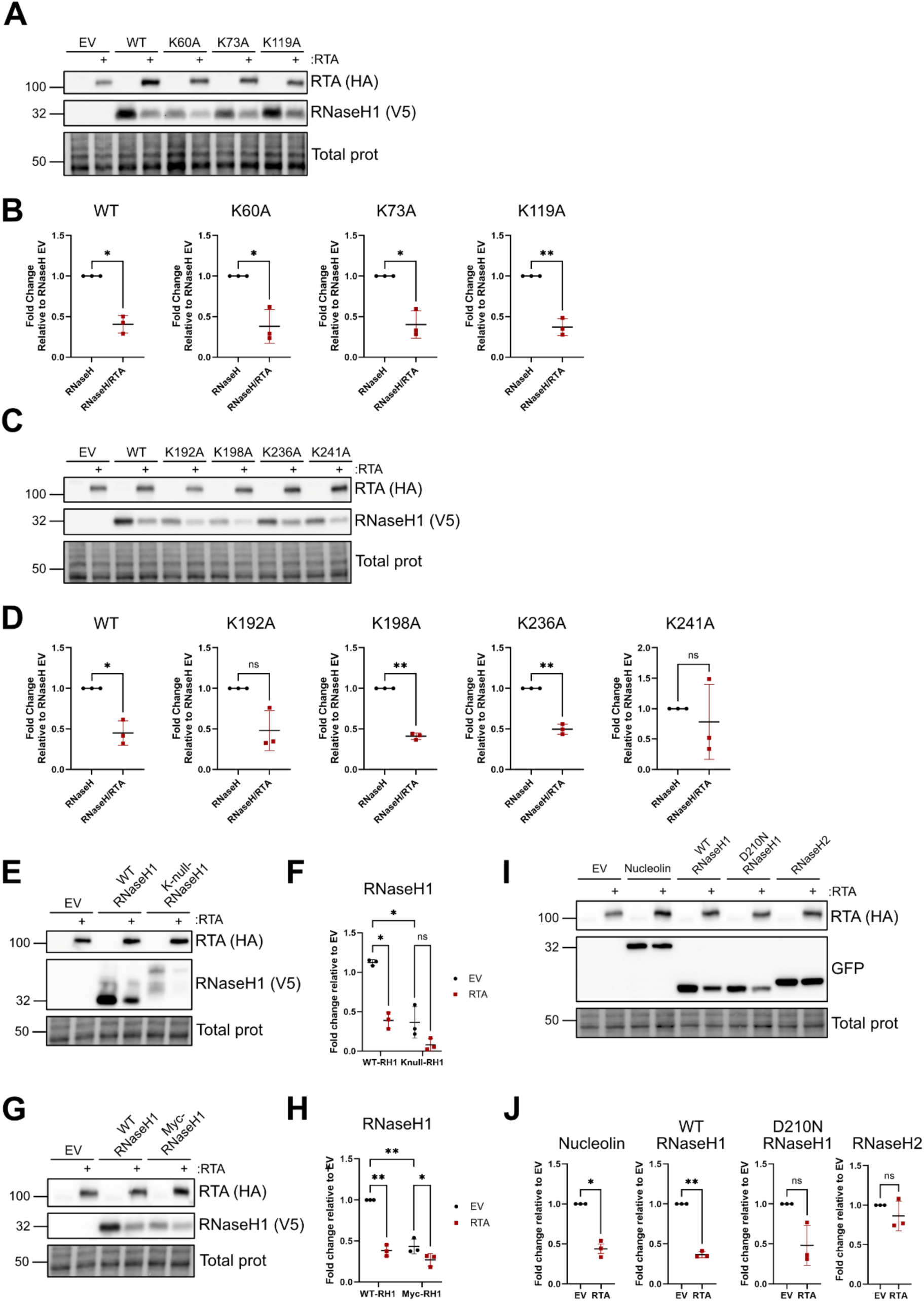
RTA does not require individual lysine residues to mediate decreased RNaseH1 protein levels. (A-D) HEK293T cells were transfected with a series of RNaseH1-expressing vectors with individual lysine to alanine mutations (K->A) in combination with an RTA-expressing vector or an empty vector (EV) control. 24 hrs post transfection cells were lysed and immunoblotting was performed using RNaseH1 (V5) and RTA (HA) specific antibodies. Samples were quantified by normalizing RNaseH1 protein levels to the total protein (total prot) in each lane using ImageLab (BioRad) and then to the respective EV co-transfection control, aka the specific RNaseH1 construct alone. Statistics were performed using a paired Student’s t-test. N=3; mean ± SD. (E-F) HEK293T cells were transfected with WT RNaseH1 or K-null RNaseH1-expressing vectors in combination with an RTA-expressing vector or an empty vector (EV) control. 24 hrs post transfection cells were lysed and immunoblotting was performed using RNaseH1 (V5) and RTA (HA) specific antibodies. Samples were quantified by normalizing RNaseH1 protein levels to the total protein (total prot) in each lane using ImageLab (BioRad) and then to the EV co-transfection control, aka WT RNaseH1 alone. Statistics were performed using a two-way ANOVA with a Tukey’s post hoc test. N=3; mean ± SD. (G-H) HEK293T cells were transfected with WT RNaseH1 or N-terminally myc-tagged RNaseH1-expressing vectors in combination with an RTA-expressing vector or an empty vector (EV) control. 24 hrs post transfection cells were lysed and immunoblotting was performed using RNaseH1 (V5) and RTA (HA) specific antibodies. Samples were quantified by normalizing RNaseH1 protein levels to the total protein (total prot) in each lane using ImageLab (BioRad) and then to the EV co-transfection control, aka WT RNaseH1 alone. Statistics were performed using a two-way ANOVA with a Tukey’s post hoc test. N=3; mean ± SD. **(I-J)** HEK293T cells were transfected with a series of GFP-tagged constructs including nucleolin, WT RNaseH1, catalytically dead RNaseH1 (D210N RNaseH1), or RNaseH2-expressing vectors in combination with an RTA-expressing vector or an empty vector (EV) control. 24 hrs post transfection cells were lysed and immunoblotting was performed using RNaseH1 (V5) and RTA (HA) specific antibodies. Samples were quantified by normalizing GFP-tagged protein levels to the total protein (total prot) in each lane using ImageLab (BioRad) and then to the EV co-transfection control, aka GFP-tagged construct alone. Statistics were performed using a two-way ANOVA with a Student’s t-test. N=3; mean ± SD. For panels A, C, E, I, and G, one representative experiment of three is shown.

We observed that K-null RNaseH1 exhibited altered migration compared to WT RNaseH1, likely due to an altered charge due to abundant K-to-A mutations. Despite this, co-expression of K-null RNaseH1 with RTA still led to a decrease in K-null RNaseH1 protein levels (Figure 9E,F). These data suggest that no single lysine or combination of lysine residues are required for RTA-mediated RNaseH1 protein decreases. In addition to lysine ubiquitination of internal residues, the N-terminus of a protein can also be ubiquitinated to mark it for degradation^20^. Recent work has shown that RTA induces the N-terminal ubiquitination of the cellular transcription factor ID2 to induce its degradation and that this was abrogated by the addition of an ID2 N-terminal tag^75^. To test if RTA might likewise target RNaseH1 for degradation by ubiquitinating its N-terminus, we constructed a dual tagged RNaseH1 construct, with a C-terminal V5 tag and an additional N-terminal myc tag. We co-transfected myc-RNaseH1-V5 with RTA or an EV control. By immunoblot analysis we found that i) the dual tagged RNaseH1 was not expressed as well as C-terminally tagged RNaseH1 and ii) the dual tagged RNaseH1 protein levels did decreased when co-expressed with RTA; however, this decrease was reduced compared to the C-terminally tagged RNaseH1 (Figure 9G,H). Although this suggests that RTA may N-terminally ubiquitinate RNaseH1 leading to its decrease, since our goal was to create an overexpression construct, the dual tagged RNaseH1 was not an ideal choice in this regard. Since our attempts to create any version of RNaseH1 that could both resist RTA-mediated decreases and be overexpressed to a high level were unsuccessful, we next asked whether all proteins decrease when co-expressed with RTA. To test this, we transfected a series of N-terminally GFP-tagged constructs including nucleolin (as an irrelevant nuclear protein control), RNaseH2, WT RNaseH1, and catalytically dead RNaseH1. Both WT and catalytically dead RNaseH1 should act as positive controls in this assay, as both have previously exhibited RTA-mediated protein decreases. We found that nucleolin, WT RNaseH1, and catalytically dead RNaseH1 were all decreased after co-expression with RTA, relative to the EV co-expression control; however, RNaseH2 was resistant to RTA-mediated decreases (Figure 9I,J). The decrease in nucleolin levels during RTA co-expression was unexpected; however, despite exhibiting a similar level of expression, RNaseH2 levels were not decreased, suggesting that RTA-mediated protein decreases are indeed protein-specific. These data suggest that RNaseH2 may serve as a viable candidate for R-loop resolution during KSHV infection due to its resistance to RTA mediated degradation.

KSHV RTA-mediated targeting of the R-loop resolver RNaseH1 may be a conserved mechanism to ensure OriLyt R-loop persistence. How the OriLyt R-loops form and are maintained during EBV and HCMV replication is unknown. Other members of the γ-herpesvirus subfamily, rhesus macaque rhadinovirus (RRV), murine gammaherpesvirus 68 (MHV-68), and Epstein Barr virus (EBV) encode homologous proteins to KSHV RTA. KSHV-RTA and MHV-68-RTA have confirmed E3 ligase activity and EBV-RTA and RRV-RTA have been shown to induce protein degradation in co-expression assays but have not been directly shown to have E3 ligase activity^25,78–80^. As multiple herpesviruses form R-loops, and multiple herpesviruses encode RTAs with either putative or confirmed E3 ligase activity, we hypothesized that the RTA-mediated degradation of RNaseH1 might be a conserved mechanism to ensure R-loop persistence in herpesvirus genomes. Using a panel of FLAG-tagged KSHV-, EBV-, MHV-68-, and RRV-RTA constructs from the Toth lab^75^, we tested whether RTA homologs from related γ-herpesviruses decreased RNaseH1 protein levels. Via immunoblot analysis we found that, relative to the EV co-expression control, co-expression of KSHV-RTA, EBV-RTA, and MHV-68-RTA led to a significant decrease in RNaseH1 protein level, whereas the co-expression of RRV-RTA did not (Figure 10).

**Figure 10.**
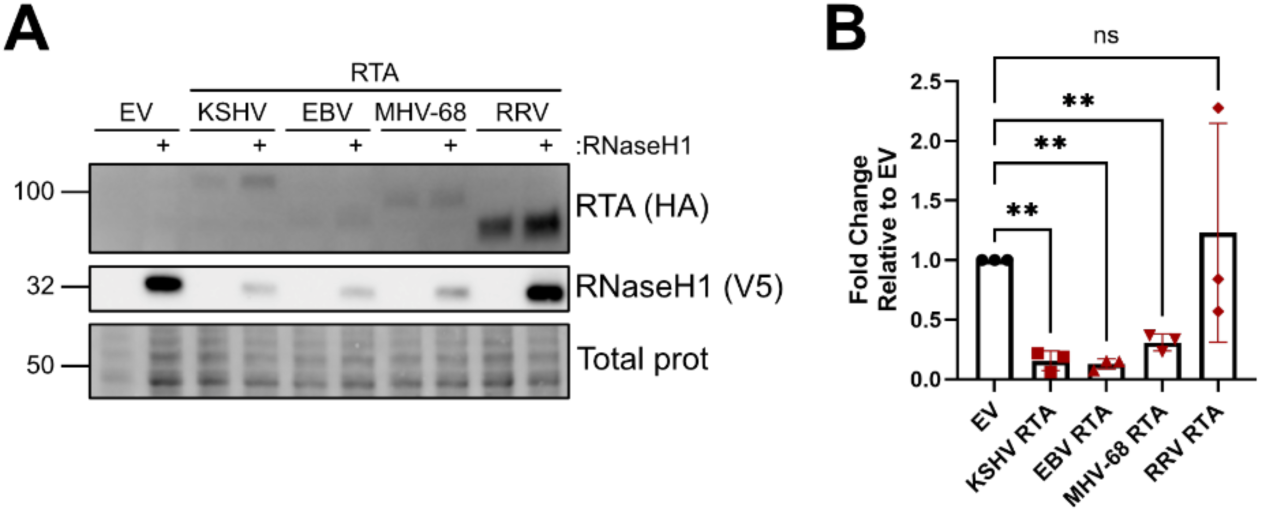
RTA homologs from related herpesviruses also decrease RNaseH protein expression. (A-B) HEK293T cells were transfected with a series of RNaseH1-expressing vectors in combination with a series of RTA-expressing vectors from different herpesviruses or an empty vector (EV) control. 24 hrs post transfection cells were lysed and immunoblotting was performed using RNaseH1 (V5) and RTA (HA) specific antibodies. Samples were quantified by normalizing RNaseH1 protein levels to the total protein (total prot) in each lane using ImageLab (BioRad) and then to the respective EV control. Statistics were performed using a one-way ANOVA with a Dunnett’s post hoc test. N=3; mean ± SD. For panel A one representative experiment of three is shown.

Given that RRV is very closely related to KSHV, and these RTA homologs have high sequence homology in the N-terminus where the E3 ligase activity of KSHV RTA is contained, this result was surprising^81–84^.

However, our findings mirror that of Combs et al. who found that KSHV-RTA, EBV-RTA, and MHV-68-RTA, but not RRV-RTA can degrade the cellular transcription factor ID2^75^. Overall, these data suggest that i) RNaseH1 is a conserved target of KSHV-RTA, EBV-RTA, and MHV-68-RTA and ii) RRV RTA acts distinctly from other RTA homologs to decrease steady-state protein levels.

## DISCUSSION

Herpesvirus genomes are predicted to contain a high number of R-loop forming sequences and both HCMV and EBV have been confirmed to form R-loops at their origins of lytic replication^63,64^. Despite this, our understanding of R-loop function during herpesvirus infection remains minimal, as does our understanding of how these structures evade the cellular machinery dedicated to R-loop resolution. Our attempts to resolve viral R-loops during KSHV infection revealed that the viral lytic switch protein, RTA, decreases the protein level of R-loop resolver RNaseH1. This suggests that viral R-loop persistence may benefit to viral replication. We show that the E3 ligase activity of RTA is at least partially required to decrease cellular RNaseH1 protein and that the RTA SIM domains are required for this effect. Finally, we find that RTA homologs from related herpesviruses MHV-68 and EBV can likewise induce a decrease in RNaseH1 protein, suggesting this mechanism is evolutionarily conserved. Taken together these data support the emerging theme of R-loop formation in herpesvirus genomes and suggest that the cellular R-loop resolver RNaseH1 may be a novel herpesvirus restriction factor.

### Are KSHV OriLyt R-loops required for viral DNA replication?

Active KSHV DNA replication occurs at one of two origins of lytic replication, OriLyt-L or OriLyt-R. Each origin contains four *cis* sequences required for replication including i) an AT-rich region, ii) a series of C/EBP binding sites, iii) an RTA-responsive element, and iv) a GC-rich OriLyt-associated transcript. In addition, active DNA replication requires eight viral proteins *in trans*, RTA, K8α, and the six core herpesvirus replication proteins including the single stranded DNA binding protein (ssDBP [ORF6]), viral DNA polymerase (ORF9), primase-associated factor (ORF40/41), helicase (ORF44), primase (ORF56) and polymerase processivity factor (ORF59)^8,73^. Together RTA and K8α associate with the core proteins to form the viral pre-replication complex that once bound to the OriLyts recruits accessory factors and initiates viral DNA replication^85^. This process is well established to occur during lytic reactivation, but it also reported to occur during primary infection to increase genome copy number prior to latency establishment^7^. Previously we found that kaposin recombinant viruses which lack the GC-rich region of the kaposin DRs have decreased intracellular genome copy number following primary KSHV infection^70^. Genomic GC-rich regions are strongly associated with R-loop formation, and R-loop formation is required for optimal viral DNA replication during EBV and HCMV infection leading us to hypothesize an analogous role for R-loops during KSHV infection^63,64^. However, our investigation of this hypothesis was confounded by several variables, discussed below, making it difficult to draw concrete conclusions.

Kaposin recombinant viruses lack R-loop formation at OriLyt-R but are still able to undergo genome copying during reactivation suggesting R-loop formation is not required for KSHV DNA replication after reactivation^70^. However, we must consider whether *T1.4*/OriLyt-L R-loop formation, which remains intact during kaposin recombinant virus replication, may act in compensatory manner to sustain DNA replication. If R-loop formation at OriLyt-L alone is sufficient to initiate genome copying during lytic reactivation this would mask our ability to detect a defect from the loss of R-loop formation at OriLyt-R. In addition, the iSLK cell lines that we used to measure kaposin recombinant virus DNA replication are estimated to maintain 50-100 copies of the viral genome^86^. The high genome copy number of iSLKs substantially increases the odds of successfully initiating genome replication, again masking our ability to defect small defects that might arise from the loss of R-loop formation at one OriLyt but not the other. In contrast to the lytic reactivation of iSLKs, during primar*y* KSHV infection there is only one incoming copy of viral DNA. This means that primar*y* infection likely represents a more stringent scenario for viral DNA replication that may result in a larger dynamic range allow which allows for subtle differences in genome copying to be resolved. In support of this, others have shown that that the transcription of the OriLyt-L associated RNA, *T1.4*, is required for viral DNA amplification specifically during primary infection but not reactivation^13^. Unfortunately, since the primary infection model for KSHV infection is inefficient, it is impractical to perform high sensitivity techniques such as ChIP-qPCR. As a result, we were unable to directly investigate R-loop function during this phase and were forced to investigate R-loops during lytic iSLK reactivation instead. It is likely that R-loop formation is required for optimal KSHV DNA replication after primary infection. However, to effectively test this, our field must either i) overcome the sensitivity issues of the primary infection model with regards to ChIP analysis or ii) resolve R-loops at both OriLyts simultaneously.

### How might KSHV OriLyt R-loops promote DNA replication?

If we assume that R-loop formation at the KSHV OriLyts is required for viral DNA replication, how might occur mechanistically? For both EBV and HCMV, defective viral DNA replication in the absence of R-loop formation is attributed to either a lack of viral ssDBP recruitment (EBV) or expression (HCMV)^63,64^. In an R-loop, the displaced single strand of DNA would provide a substrate for viral ssDBP binding, so it follows that a lack of R-loop formation might lead to decreased ssDBP recruitment. Prior to this work, the role of R-loop formation in the recruitment of viral or cellular proteins to the KSHV OriLyts was unknown. Since kaposin recombinant viruses exhibit R-loop formation at one OriLyt (OriLyt-L) but not the other (OriLyt-R), this placed us in a unique position to evaluate this question for three viral proteins, RTA, K8α, and ORF6. Using ChIP-qPCR, we found that RTA bound with comparable efficiency to OriLyt-R during WT, ΔKapABC, and ΔKapB virus infection, indicating that R-loop formation is not required for RTA recruitment. In contrast, we found that during ΔKapABC but not WT reactivation there was a slight increase in K8α binding to the OriLyt-L compared to OriLyt-R; this suggests that the transcription of a shorter transcript with less sites for K8α binding does impair K8α recruitment^13^. Finally, we found no statistically significant difference in ssDBP (ORF6) recruitment to OriLyt-R in the absence of R-loop formation. These data suggest that R-loop formation only plays a minor role in the recruitment of viral proteins to the OriLyt-R. Whether there is an R-loop dependent decrease in ORF6 protein expression during kaposin recombinant virus replication, analogous to HCMV, should be tested in future work.

Alternatively, it is possible the role of R-loop formation during KSHV reactivation may be independent of ORF6 binding and/or expression. For example, R-loops are known to recruit numerous host proteins including DNA repair factors such as Cockayne syndrome B (CSB), RAD52, breast cancer type 2 susceptibility protein (BRCA2), Poly [ADP-ribose] polymerase 1 (PARP1), and replication protein A (RPA) and R-loop formation has been shown to activate both ataxia telangiectasia mutated (ATM) and ataxia telangiectasia and Rad3-related protein (ATR)^47,48,87–93^. Herpesvirus R-loop formation could serve to localize DNA damage response (DDR) proteins to the site of the viral genome replication that are subsequently needed for viral DNA replication. In support of this idea, both meiotic recombination 11 (MRE11) and replication protein A 34 kDa subunit (RPA32) have been shown to localize to both KSHV and EBV replication compartments and during EBV infection the knockdown of RPA32 prevents viral DNA replication^94,95^. PARP1 has also been shown to be recruited the KSHV OriLyts and pharmacological inhibition of PARP1 activity decreases viral DNA replication^96^. Finally, activation of ATM and subsequent phosphorylation of Sp1 has also been shown to be required for EBV DNA replication^97^. Further work is required to determine whether DDR proteins localize to the KSHV genome in an R-loop dependent manner. A potential model for how R-loop formation may benefit viral replication is shown in Figure 11.

**Figure 11.**
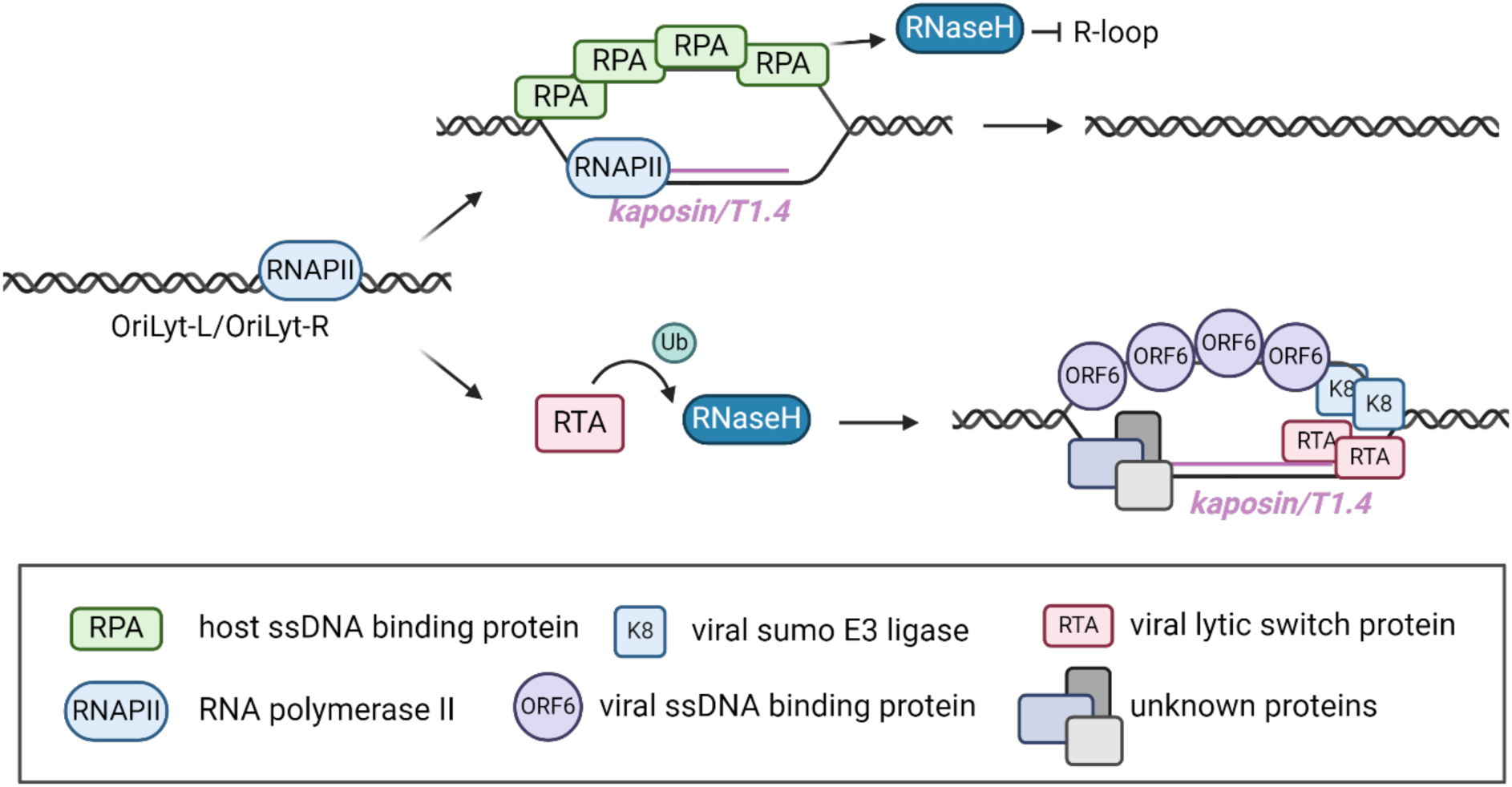
Proposed model for how R-loop formation at the KSHV OriLyts may benefit viral replication. A graphical depiction of how R-loop formation could benefit viral replication is shown. Briefly, transcription of *T1.4* and *kaposin* at OriLyt-L and OriLyt-R, respectively, leads to R-loop formation. Normally this structure would be resolved by cellular RNaseH1 (top panel) however during KSHV infection RTA targets RNaseH1, decreasing its levels. Decreased RNaseH1 leads to R-loop persistence which could allow for viral proteins such as K8α and ORF6, or unknown proteins such as DNA repair factors, to be recruited. These proteins are required for subsequent viral DNA replication.

### How do the E3 ligase and SIM domains of RTA contribute to RNaseH1 decreases?

KSHV RTA targets several proteins for degradation either directly or by cooperating with cellular E3 ligases ITCH and UBE3C^76,77^. As an E3-like ligase, RTA has an N-terminal Cys/His rich domain and undergoes autoubiquitination and degradation^25,27^. Here we showed that KSHV reactivation led to a decrease in RNaseH1 protein level, and that the overexpression of RTA alone was sufficient for this effect. To test whether RTA uses E3 ligase activity to target RNaseH1 we created an RTA construct with a mutated E3 catalytic domain. E3-ligase dead RTA expressed better than WT, suggesting that our mutation of C141S and H145L was effective at decreasing E3 ligase activity. We found expression of this construct led to a small rescue in RNaseH1 protein levels, however a partial decrease still occurred.

These results reveal that RTA E3 ligase activity is at least partially required for decreased RNaseH1 levels during co-expression; however, the point mutations we introduced may not completely abolish this activity, leading to an incomplete rescue of RNaseH1 levels. These data may also be explained by the possibility that RTA recruits a host E3 ligase to assist with RNaseH1 ubiquitination and this putative interaction is unperturbed by our mutagenesis. We tested the two host E3 ligases previously identified to act in concert with RTA, ITCH and UBE3C, and found that their knockdown did not alter RTA-mediated RNaseH1 decreases. We were unable to assess the efficiency of this knockdown via immunoblot, therefore the decrease we observed in ITCH and UBE3C transcript levels may not correspond to a functional protein knockdown. Immunoblotting using ITCH and UBE3C specific antibodies should be performed before these results can be fully interpreted. In addition, RTA may interact with a yet-to-be identified host E3 ligase to promote RNaseH1 degradation which we did not assess. RTA contains multiple SIM domains and has SUMO-targeting E3 ligase activity (STUbL) activity^28^. We found that, unlike E3 ligase dead RTA, co-expression of SIM-mutant RTA completely abolished RNaseH1 decreases. This reveals that the ability of RTA to interact with SUMO moieties is of critical importance to its ability to promote decreased RNaseH1 levels. These data may be explained in two ways: i) WT RTA interacts directly with a SUMO-modification on RNaseH1 to directly promote its degradation or ii) WT RTA interacts with an unidentified protein via its SIM domains and this protein regulates RTA E3 ligase activity in an unknown manner. In support of the latter hypothesis, during overexpression we observed that SUMO-mutant RTA exhibited an altered immunoblot banding pattern that included multiple faster migrating bands, suggesting SIM-mutant RTA fails to undergo certain PTMs which could affect its E3 ligase activity. For example, if by virtue of its inability to interact with SUMO-modified proteins, SIM-mutant RTA does not gain a specific PTM it may be improperly localized to the cytoplasm where it would be rendered unable to target RNaseH1 for degradation. Whether RTA acts directly via SUMO-modified RNaseH1 or indirectly via SUMO-mediated interactions with other proteins will be the subject of future studies.

### How is RNaseH1 marked for protein turnover by RTA?

To measure the effect of OriLyt R-loop formation on viral DNA replication, we tried to overexpress the cellular R-loop resolver RNaseH1 but found it to be rapidly decreased by RTA in a partially E3 ligase-dependent manner. E3 ligase-mediated ubiquitination of internal lysine residues marks proteins for autophagic or proteasomal degradation. In the hopes of generating an RNaseH1 construct that is resistant to RTA-mediated decreases, we created a series of lysine mutated RNaseH1 constructs. By testing these constructs in our co-expression model, we found that none of the lysine residues previously predicted to undergo ubiquitination were singularly required for RTA-mediated RNaseH1 decreases. Moreover, a completely lysine-null RNaseH1 still exhibited decreased protein levels during RTA co-expression, albeit to a lesser extent. These data suggest that RTA is not solely dependent on internal lysine residues to mediate RNaseH1 targeting. RTA has previously been shown to target the N-terminus of a cellular transcription factor for ubiquitination and subsequent degradation, an effect which could be abrogated through the addition of an N-terminal FLAG tag^20,75^. We found that the addition of an N-terminal myc tag partially protected RNaseH1 from RTA-mediated degradation. Given that E3 ligase activity is at least partially required for RTA-mediated RNaseH1 decreases, we envision three possible explanations for these data. First, RTA uses a combination of both internal lysine residues and N-terminal ubiquitination to mediate RNaseH1 decreases. Second, RTA induces RNaseH1 degradation by ubiquitinating an internal non-lysine residue such as a cysteine, serine, or threonine via thioester (cysteine) or oxyester bonds (serine and threonine)^18,98–100^. Third, a double negative scenario occurs in which RTA ubiquitinates a negative regulator of ubiquitin-independent RNaseH1 degradation. For example, the phosphorylation of Tau by non-proline-directed Ca^2+^/calmodulin-dependent protein kinase II (CaMKII) inhibits its direct degradation by the 20*S* proteasome^101^. If RTA degrades a negative regulator, this could promote the degradation of RNaseH1. Future work will determine the exact mechanism of KSHV RTA-mediated RNaseH1 degradation.

### Significance

Here we show that like other herpesviruses, R-loops form at the KSHV OriLyts. To determine the function of OriLyt-associated R-loop formation during KSHV infection, we tried to resolve them by overexpressing RNaseH1. Yet our attempts to do so were thwarted by RTA because its expression led to decreased RNaseH1 levels. This finding suggests KSHV uses RTA to promote R-loop persistence and may explain how KSHV OriLyt-associated R-loops evade resolution. RTA homologs from other у-herpesviruses likewise decrease RNaseH1 levels suggesting RNaseH1 may be a novel herpesvirus restriction factor.

While an unexpected and exciting finding, this rendered us unable to resolve R-loops as planned and thus we have not yet been able to determine their function in the context of KSHV infection. In preliminary work, we observed protein levels of RNaseH2 were not decreased by the presence of RTA co-expression, suggesting future experiments on RNaseH2 overexpression for R-loop resolution during infection.

### Limitations

i) While RTA is sufficient for RNaseH1 protein decreases and its E3 ligase activity is partially required for this event, we did not directly demonstrate that RTA leads to an increase in RNaseH1 ubiquitination. ii) For RTA to ubiquitinate RNaseH1 directly, RTA and RNaseH1 must interact, but we have not yet validated this interaction. iii) Despite numerous attempts, we were unable to create an RTA-resistant RNaseH1 construct and therefore unable to directly test whether RNaseH1-mediated R-loop resolution decreases KSHV DNA replication. We aim to overexpress the R-loop resolver such as RNaseH2 to overcome this barrier.

## METHODS

### Cell culture

All cells were grown at 37°C with 5% CO_2_ and atmospheric O_2_. HEK293T cells (ATCC) were cultured in Dulbecco modified Eagle medium (DMEM; Thermo Fisher) supplemented with 100 U/mL penicillin, 100 μg/mL streptomycin, 2 mM l-glutamine (Thermo Fisher), and 10% fetal bovine serum (FBS; Thermo Fisher). iSLK.TREx-RTA cells were cultured likewise with the addition of 1 μg/mL puromycin and 250 μg/mL Geneticin (Thermo Fisher). BAC16-iSLK.RTA cells were cultured similar to iSLK.TREx-RTA cells, with the addition of 1200 μg/mL hygromycin B (Thermo Fisher). BCBL1-TREx-RTA cells were cultured in RPMI 1640 (ThermoFisher) supplemented with 55 μM β-mercaptoethanol, 100 U/mL penicillin, 100 µg/mL streptomycin and 10% FBS.

### Drug treatments

MG132 (Sigma-Aldrich) and CQ (Sigma-Aldrich) were dissolved in either DMSO or nuclease free water, respectively, and added to a small volume of spent media prior to being spiked in to each well individually at a final concentration of 10 µM and 25 µM, respectively, for a duration of four hours.

### Site-directed mutagenesis

Site-directed mutagenesis PCR was performed was performed using the primers listed in Table 1, the forward primer of which was always phosphorylated. Briefly, these primers were used to amplify the entire plasmid thereby incorporating the desired mutation. The template plasmid was then digested using DpnI (NEB) and the resulting product was ligated, transformed, prepped, and sequenced prior to use.

**Table 1.**
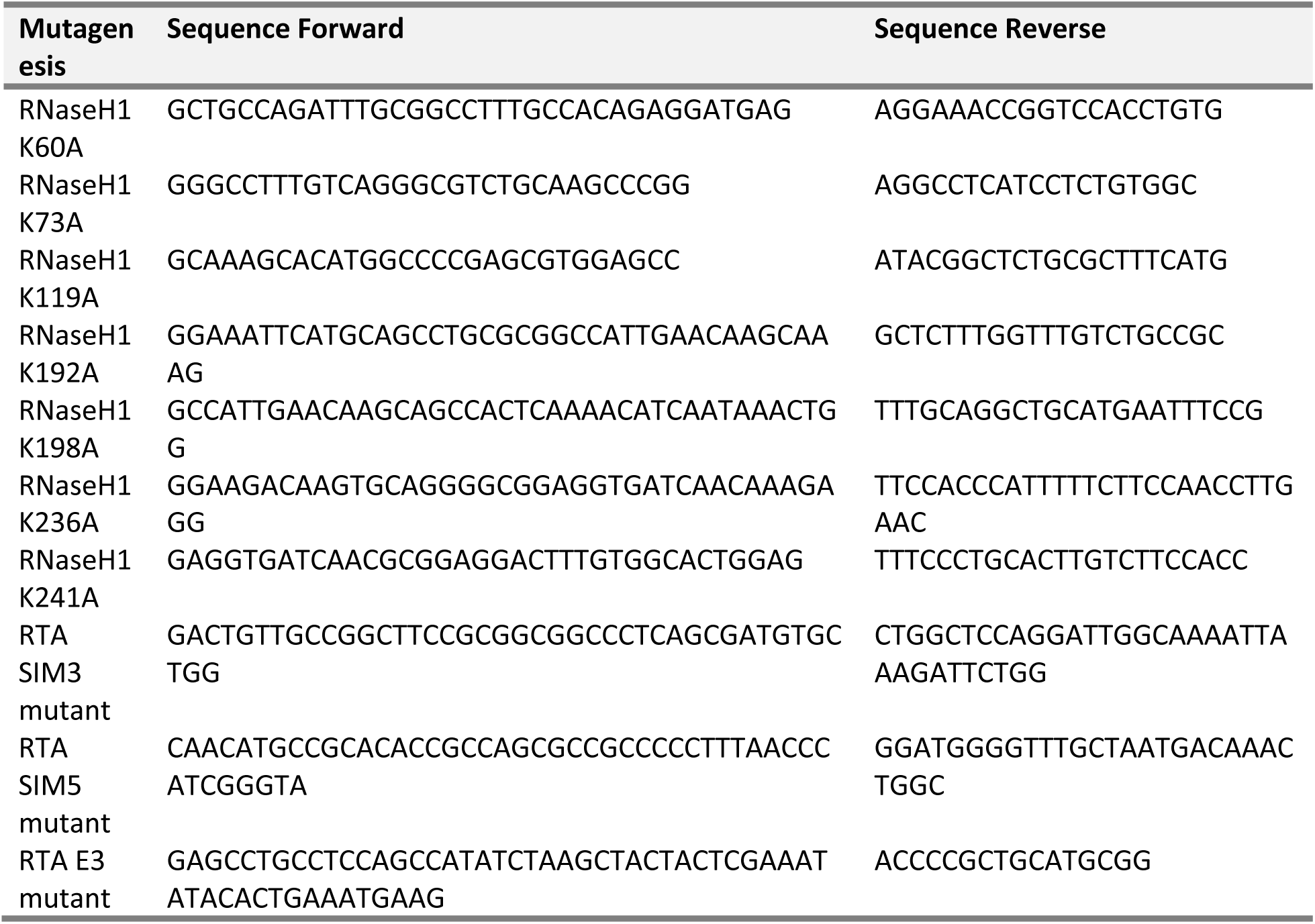
Site directed mutagenesis primers.

### Transfection

HEK293T cells seeded to 70% confluency in antibiotic-free 10% FBS–DMEM were washed once with PBS, and medium was replaced with 800 uL of serum-free, antibiotic-free DMEM per well prior to transfection. Cells were transfected with plasmids of interest (Table 2) using polyethylenimine (PEI, Polysciences). At 4 h post-transfection, the medium was replaced with antibiotic-free 10% FBS–DMEM.

**Table 2.**
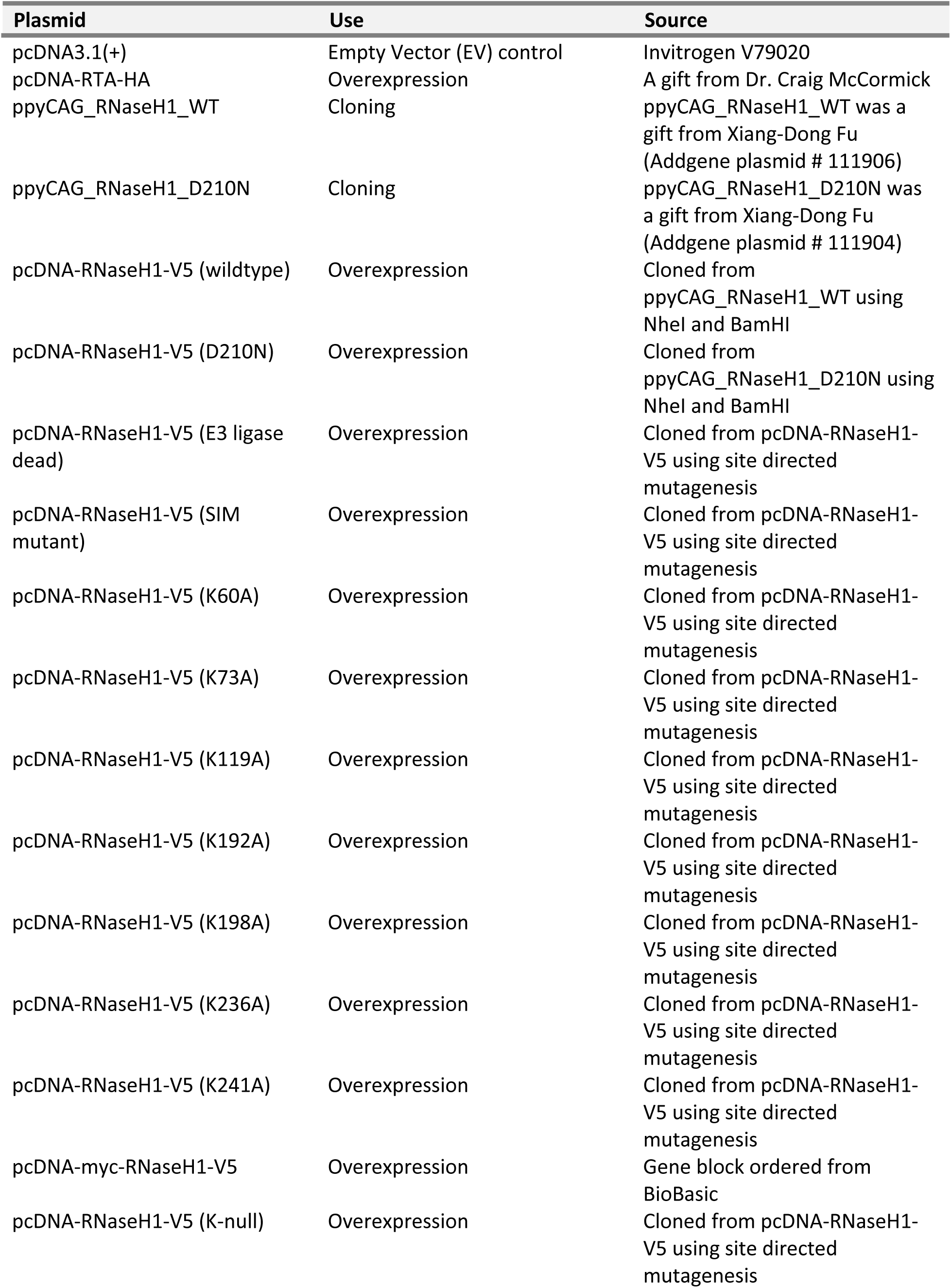
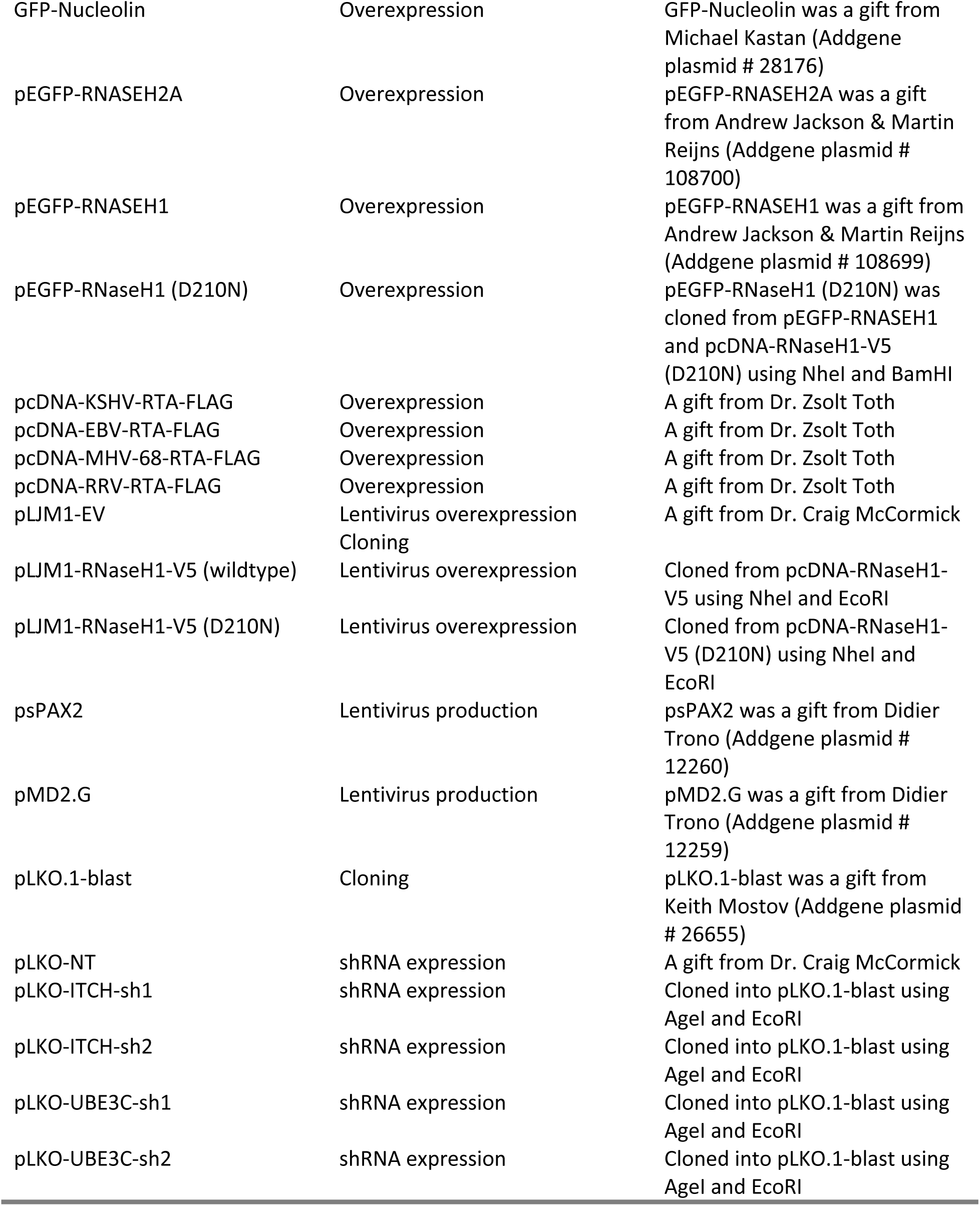
Plasmids.

### RT-qPCR

RNA was collected using either the RNeasy Plus Mini Kit (Qiagen) or TRIzol extraction according to the manufacturer’s instructions and stored at −80°C until further use. RNA concentration was determined using NanoDrop OneC (ThermoFisher) and 500 ng of RNA was reverse transcribed using MaximaH (ThermoFisher) with a combination of random hexamer and oligo dT primers, according to the manufacturer’s instructions. For qPCR cDNA was diluted 1in10 and SsoFast EvaGreen Mastermix (Biorad) was used for amplification. The ΔΔquantitation cycle (Cq) method was used to determine the fold change in expression of target transcripts using 18S as a house-keeping control gene. Primer sequences for bqPCR can be found in Table 3.

**Table 3.**
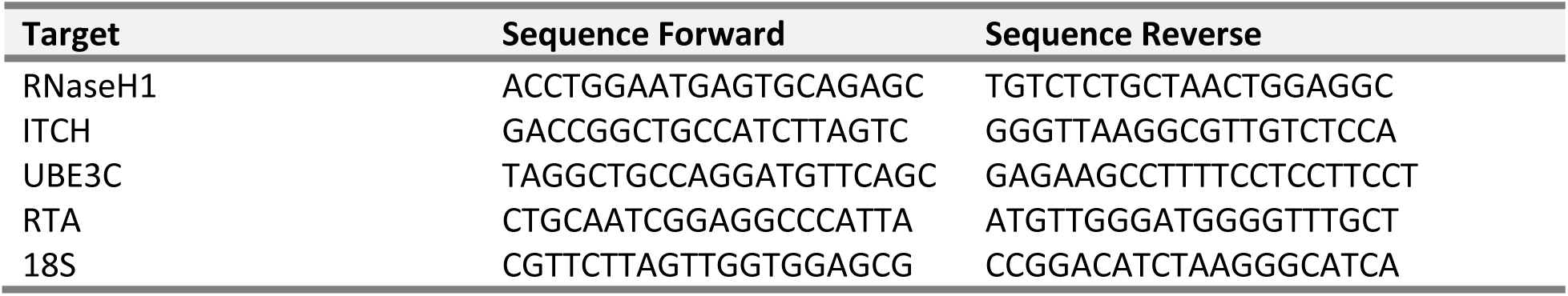
qPCR primers.

### Lentivirus production and transduction

All recombinant lentiviruses were generated using a second-generation system. HEK293T cells were transfected with psPAX2, MD2-G, and the lentiviral transfer plasmid containing either a gene interest (pLJM1) or shRNA (pLKO) using polyethylenimine (PEI, Polysciences). 4 h post-transfection, serum-free media was replaced with DMEM containing serum but no antibiotics. Viral supernatants were harvested 48 h post-transfection and stored at −80°C prior to use. For transduction, lentiviruses were thawed at 37°C and added to target cells in complete media containing 5 μg/mL polybrene (Sigma-Aldrich). After 24 hours, the media was replaced with selection media containing 5 μg/mL blasticidin (ThermoFisher) and cells were selected for 48 h before proceeding with experiments. Targeting sequences for shRNA knockdown can be found in Table 4.

**Table 4.**
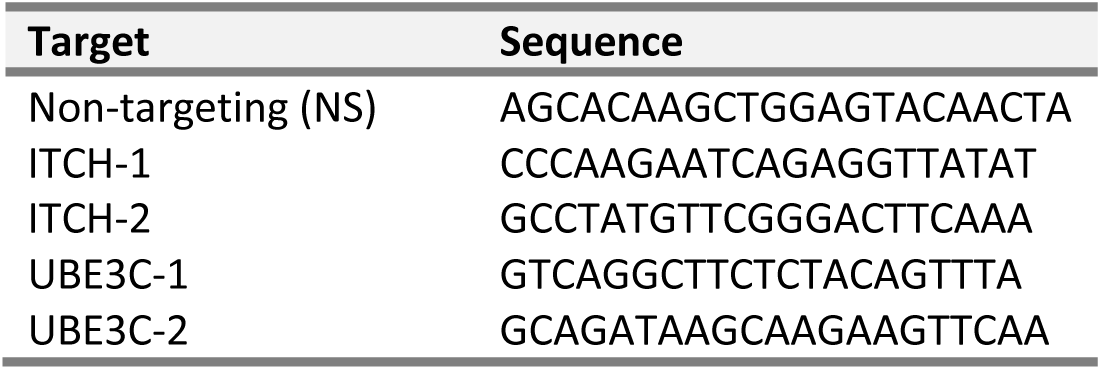
shRNA sequences.

### Chromatin Immunoprecipitation (CHIP) qPCR

15 cm dishes of latent BAC16-iSLK cells were grown to 90% confluency, formaldehyde was added to a final concentration of 1%, and cells were cross-linked for 10 mins at 37°C. Glycine was added to a final concentration of 125 mM for 10 min at RT to quench the formaldehyde. Media was removed and cells were washed with 10 mL PBS prior to being harvested by scraping into 1 mL PBS. Cells were pelleted at 800 x g for 5 mins and supernatant was removed. Pellets were snap frozen using liquid N_2_ and stored at - 80°C for no less than a month prior to use. To isolate the chromatin for immunoprecipitation, pellets were thawed on ice, resuspended in freshly prepared 600 μL ChIP lysis buffer (50 mM HEPES [pH 7.5], 140 mM NaCl, 1 mM EDTA [pH 8.0], 1% Triton-X-100, 0.1% sodium deoxycholate, 0.1% SDS, protease inhibitor) and sonicated in a water bath sonicator (Bioruptor™ UCD-200, Diagenode) for 1 h (30 s on, 90 s off intervals). Sonicated chromatin was clarified by centrifugation at 8000 x g for 10 min at 4°C and 200 μL of sonicate per IP was added to 800 μL freshly prepared RIPA buffer (50 mM Tris-HCl [pH 8.0], 150 mM NaCl, 2 mM EDTA [pH 8.0], 1% NP-40, 0.5% sodium deoxycholate, 0.1% SDS, protease inhibitor), 50 uL of the diluted sonicate was removed and stored at −20°C for an input control. Experimental or IgG control antibodies (Table 6) were added and samples were incubated while rotating at 4°C for 1 h.

Meanwhile, 25 μL of SureBeads™ Protein A magnetic beads (BioRad) per IP were washed 3X in 1 mL RIPA buffer and then resuspended in 500 μL RIPA buffer with 75ng/μL ssDNA (ThermoFisher) and 0.1 μg/μL BSA (Sigma-Aldrich) and incubated for 30 min at RT to block. Post bock, beads were washed 1X with 1 mL RIPA buffer, resuspended in 100 μL RIPA buffer per IP, added to the sonicate/antibody samples and left to incubate while rotating at 4°C overnight. The following day antibody/bead complexes were pelleted using a mag rack (BioRad), supernatant was removed, and the beads were sequentially washed with 1 mL freshly prepared Low Salt Wash Buffer (0.1% SDS, 1% Triton-X-100, 2 mM EDTA [pH 8.0], 20 mM Tris-HCl [pH 8.0], 150 mM NaCl), High Salt Wash Buffer (0.1% SDS, 1% Triton-X-100, 2 mM EDTA [pH 8.0], 20 mM Tris-HCl [pH 8.0], 500 mM NaCl), LiCl Wash Buffer (0.25 M LiCl, 1% NP-40, 1% sodium deoxycholate, 1 mM EDTA [pH 8.0], 10 mM Tris-HCl [pH 8.0]). Supernatant was removed, 120 μL of Elution Buffer (1% SDS, 4% sodium deoxycholate) was added, and samples were incubated at 65°C for 1 h with occasional agitation. Beads were pelleted and the elute was removed and transferred to a new tube. Simultaneously, input samples were thawed and 70 μL of RIPA buffer was added to bring the final volume to 120 μL NaCl and RNaseA (Thermo Fisher) were added to all samples to a final concentration of 0.2 M and 30 μg/μL respectively, samples were incubated at 65°C for 1 h followed by the addition of 3.2U proteinase K (Thermo Fisher) and incubation at 60°C for 1 h. Post incubation, a 1:1 volume of phenol-chloroform-isoamyl alcohol (Thermo Fisher) was added and samples were briefly vortexed and then centrifuged at 14 000 x g for 10 min at RT. The aqueous phase was removed and transferred to a new tube to which 0.1 volumes 3M sodium acetate, 2.5 volumes 100% EtOH and 20 μg glycogen (Thermo Fisher) was added and DNA was precipitated overnight at −80°C. Precipitated DNA was pelleted at 14 000 x g for 30 min at 4°C, washed with 500 μL 70% EtOH, and pelleted again at 14 000 x g for 5 min at 4°C. EtOH wash was removed, pellet was air dried, resuspended in 50 μL nuclease free water, and used directly in qPCR. Primer sequences can be Table 5.

**Table 5.**
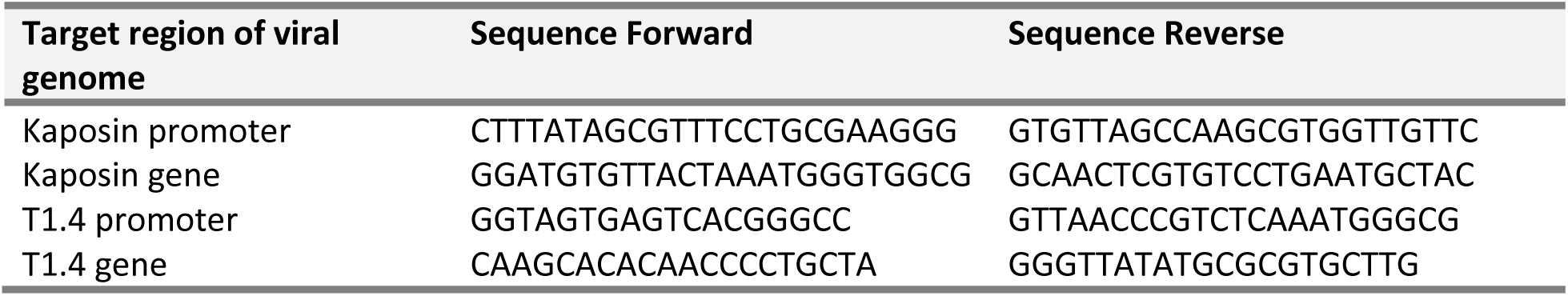
ChIP qPCR primers.

### Immunoblotting

Cells were lysed in 1× Laemmli buffer (4% SDS, 20% glycerol, 120 mM Tris-Cl [pH 6.8] and ddH_2_O) and stored at −20°C until use. The DC protein assay (Bio-Rad) was used to quantify total protein concentration as per the manufacturer’s instructions. 10 to 15 μg of protein lysate was resolved by SDS-PAGE on Tris-Glycine eXtended (TGX) Stain-Free acrylamide gels (Bio-Rad). Stain-Free technology uses a proprietary trihalo compound that binds to tryptophan residues^102^. Following a short UV-activation, trihalo-bound proteins generate a fluorescent signal that allows them to be readily visualized. According to Uniprot, 90% of proteins contain tryptophan, meaning that the resulting signal is an effective representation of most proteins present in each sample (total protein). This method is preferred over traditional “house-keeping” loading controls such as β-actin and Glyceraldehyde 3-phosphate dehydrogenase (GAPDH), the levels of which can be easily altered by experimental conditions.

Membranes were blocked in 5% bovine serum albumin (BSA) or 5% skim milk in Tris-buffered saline-Tween 20 (TBS-T). Primary and secondary antibodies were diluted in 2.5% BSA or 2.5% skim milk, dilutions can be found in Table 6. Membranes were visualized using ProtoGlow ECL (National Diagnostics) and the ChemiDoc Touch Imaging system (Bio-Rad).

**Table 6.**
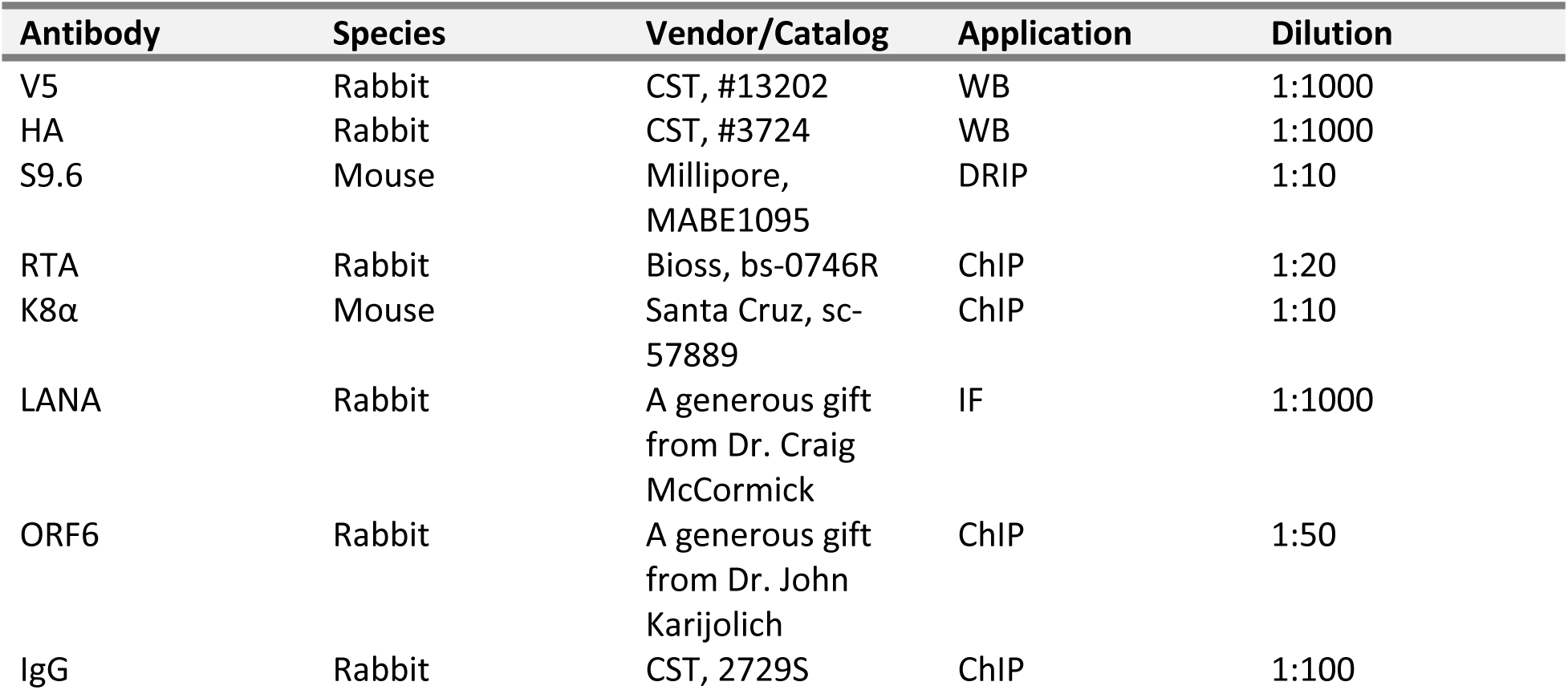
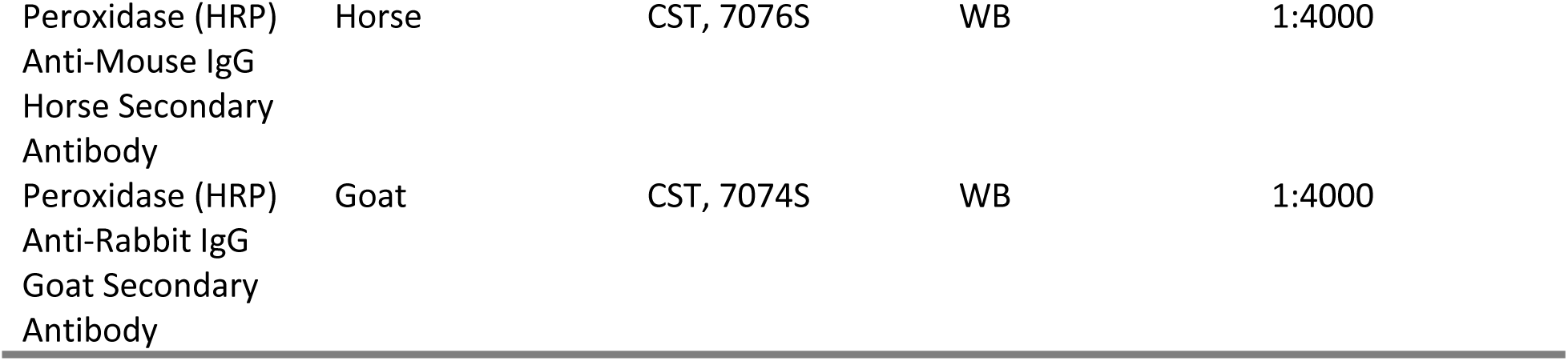
Antibodies.

### Statistics

Statistics were performed using Graphpad Prism version 9 or 10.2.2 except for those specific to RNA-seq which were performed separately. The specific statistical test is specified in the figure legend.

## Graphing

All graphing was done using Graphpad Prism version 9 or 10.2.2.

## ACKNOWLEDGEMENTS

The authors sincerely thank the members of the Corcoran lab for helpful discussions. The authors would like to thank Dr. Zsolt Toth (University of Florida) for sharing expression plasmids for other herpesvirus RTA molecules and Dr. John Karijolich (Vanderbilt University Medical Center) for the ORF6 antibody. Plasmids kindly gifted to us through Addgene are detailed in Table 3. MK was supported by a Canadian Institutes for Health Research Doctoral Graduate Scholarship and a Cumming School of Medicine Doctoral training award. Operating funds to support this work derive from a Canadian Institutes for Health Research Project Grant PJT-183595 awarded to JAC. Graphics were created using BioRender or Affinity Designer.

## Author Contributions

Mariel Kleer: Conceptualization, Experimentation, Data Analysis, Paper Writing, Paper Editing

Julia Fox: Experimentation, Paper Editing

Jennifer A. Corcoran: Conceptualization, Supervision, Funding Acquisition, Project Administration, Paper Writing, Paper Editing

